# ZNF692 organizes a hub for ribosome maturation enhancing translation in rapidly proliferating cells

**DOI:** 10.1101/2022.05.26.493655

**Authors:** M.Carmen Lafita-Navarro, Yi-Heng Hao, Chunhui Jiang, Isabella N. Brown, Seoyeon Jang, Niranjan Venkateswaran, Elizabeth Maurais, Weronika Stachera, Tsung-Cheng Chang, Dorothy Mundy, Jungsoo Han, Vanna M. Tran, Marcel Mettlen, Jeffrey B. Woodruff, Joshua T. Mendell, Nick V Grishin, Lisa Kinch, Michael Buszczak, Maralice Conacci-Sorrell

**Affiliations:** Department of Cell Biology, University of Texas Southwestern Medical Center, Dallas, Texas 75390, USA; Department of Molecular Biology, University of Texas Southwestern Medical Center, Dallas, Texas 75390, USA; Live Cell Imaging Core Facility, University of Texas Southwestern Medical Center, Dallas, Texas 75390, USA; Hamon Center for Regenerative Science and Medicine, University of Texas Southwestern Medical Center, Dallas, TX 75390, USA; Howard Hughes Medical Institute, University of Texas Southwestern Medical Center, Dallas, TX 75390, USA; Harold C. Simmons Comprehensive Cancer Center, University of Texas Southwestern Medical Center, Dallas, Texas 75390, USA; Department of Biophysics, University of Texas Southwestern Medical Center Dallas TX USA

**Keywords:** MYC, nucleolus, ribosome biogenesis, protein synthesis, ZNF692, ZFP692, NPM1, rRNA, exosome, 18S, EXOSC7, EXOSC8, KRR1, 40S

## Abstract

Rapidly proliferating cells produce more ribosomes to translate sufficient proteins for cell growth. One of the first and rate limiting steps in translation initiation is the interaction of the small ribosomal subunit with mRNAs. Therefore, effective small ribosomal subunit biogenesis is critical for translation initiation efficiency. Here we report the identification of the zinc finger protein 692 (ZNF692), a MYC-induced nucleolar scaffold that coordinates the final steps in the biogenesis of the small 40S ribosome. ZNF692 forms a complex with rRNA, the 90S processome and the nucleolar exosome in the granular component of the nucleolus creating a hub specialized in the final steps of 18S processing and small ribosomal subunit maturation. Cancer cells are more reliant on ZNF692 for increased translation than normal cells. We propose that MYC increases translation efficiency by promoting the expression of ZNF692, adjusting the translation rate to the increase in mRNA transcription induced by MYC.

---

The nucleolus is one of the largest cellular organelles, which is responsible for every step in ribosome biogenesis ^1^. Dysregulation of ribosome biogenesis is reflected by alterations in nucleolar morphology ^2, 3^ and can lead to a variety of human diseases including Alzheimer’s, Parkinson’s, Huntington’s diseases, Diamond Blackfan Anemia, and Treachers-Collins Syndrome ^1–8^. Increased nucleolar size and activity are associated with augmented ribosome production in cancer cells ^9^. Indeed, having more and larger nucleoli is the most distinguishable morphological alteration of a cancer cell and is frequently used by pathologists to grade solid tumors ^10, 11^.

The major steps necessary for ribosome biogenesis are spatially organized in three distinct nucleolar compartments: the fibrillar center (FC), the dense fibrillar component (DFC), and the granular component (GC), all named based on their morphological appearance under electron microscopy (EM). rDNA is transcribed in the nucleolus as a single precursor transcript (47S pre-rRNA) by RNA polymerase I (RNAPol I) at the border of the FC and DFC. Human cells have around 400 ribosomal DNA (rDNA) loci per diploid genome from which 20-50% are actively transcribed in the nucleolus of proliferating cells ^1, 12^. The processing of the 47S pre-rRNA starts in the FC and progresses to later steps in the DFC and GC to generate the 18S, 5.8S, and 28S ribosomal RNAs (rRNA). The 18S rRNA constitutes the 40S small ribosomal subunits and the 5.8S, 28S, and 5S, which is synthesized from a separate locus by RNA polymerase III (RNAPol III) ^13^, constitute the 60S large ribosomal subunits ^14^.

The earliest pre-ribosome, named 90S pre-ribosome or small subunit processome, is assembled co-transcriptionally on pre-rRNA ^15^. Pre-rRNA is cleaved giving rise to two precursor complexes that generate mature small 40S and large 60S ribosomal subunits, respectively ^15, 16^. Recent studies have highlighted major differences in the formation of the 3′ end of the 18S rRNA between yeast and mammals. While this process is accomplished through endonucleolytic cleavages in yeast, it requires the combined action of both endonucleases and exonucleases in human cells ^15, 17^.

The RNA exosome performs 3’-5’exoribonucleolytic activity to degrade or process RNA ^18–20^. The exosome comprises nine structural subunits that are organized in a ring structure ^18, 21^. The core subunits EXOSC4–EXOSC9 form a barrel-like structure. The subunits EXOSC1-3 form a central pore cap that binds RNA and other cofactors providing substrate specificity. Nucleolar exosomes, which associate with the 3’-5’exonuclease EXOSC10/RRP6 ^22^, are involved in the maturation steps that generate 18S and 5.8S rRNA in human cells. Depletion of EXOSC10/RRP6 causes an accumulation of 21S rRNA and a reduction of its product, the near mature 18S-E pre-rRNA ^23, 24^. The 18S-E pre-RNA complexed with ribosomal proteins is exported into the cytoplasm where it matures into 18S rRNA within the 40S small ribosome subunit ^17^. 40S can then interact with mRNA and translation initiation factors forming a complex that recruits the 60S ribosomal subunit to form actively translating ribosomes ^25^.

Numerous studies have shown that the transcription factor MYC promotes ribosome biogenesis ^26, 27^ by driving rDNA transcription, regulating the transcription of structural and regulatory components of the ribosome, and amplifying RNAPol III-mediated 5S rRNA production ^9, 28–30^. Moreover, MYC-induced transformation was found to be dependent on increased ribosome biogenesis and protein synthesis ^27^. Here, we describe the identification and characterization of ZNF692, a protein that evolved in chordates, as a MYC-induced nucleolar scaffold responsible for increasing the efficiency of small ribosomal subunit maturation in mammalian cells. However, our work shows that ZNF692 localizes in the GC of the nucleolus where it facilitates the formation of a complex that anchors the exosome complex to the 40S processome, allowing 21S rRNA trimming and thus 18S maturation. We propose that by facilitating the final steps of 40S subunit biogenesis, MYC-induced ZNF692 increases translation efficiency to match protein synthesis with the overall increase in mRNA levels that are transcribed by MYC in proliferative cells.

## Results

### MYC promotes the transcription of the novel and evolutionarily conserved nucleolar factor ZNF692

To dissect the regulatory network underlying cell growth, we performed RNA-seq experiments comparing Rat1 *myc*-/- fibroblasts expressing empty vector or reconstituted with human MYC, and searched for transcription factors with increased expression in MYC-expressing cells ^31–34^. We identified three understudied Zn finger-containing genes whose expression were upregulated by MYC (Supplementary Fig. 1A, 1B). Among these proteins, ZFP692 (ZNF692 mouse orthologue) was the only Zn finger protein identified to also be induced by MYC in previously published datasets of MYC-driven liver tumors ^35^, and lymphoma ^36^ (Supplementary Fig. 1C, D). RT-qPCR and Western blots (WB) in Rat1 fibroblasts and in human retinal pigment epithelial cells (ARPE-19 cells) confirmed the ZFP692/ZNF692 mRNA and protein induction by MYC (Fig. 1A-D). Moreover, knocking down *MYC* in MYC-reconstituted *myc-/-* fibroblasts decreased the levels of ZFP692 mRNA and protein (Fig 1A and Supplementary Fig. 1E). ChIP-seq datasets deposited in ENCODE revealed that MYC bound to the ZNF692 promoter in four independent cell lines (Supplementary Fig. 1F), demonstrating that ZNF692 is a direct and universal transcriptional target of MYC.

**Figure 1:**
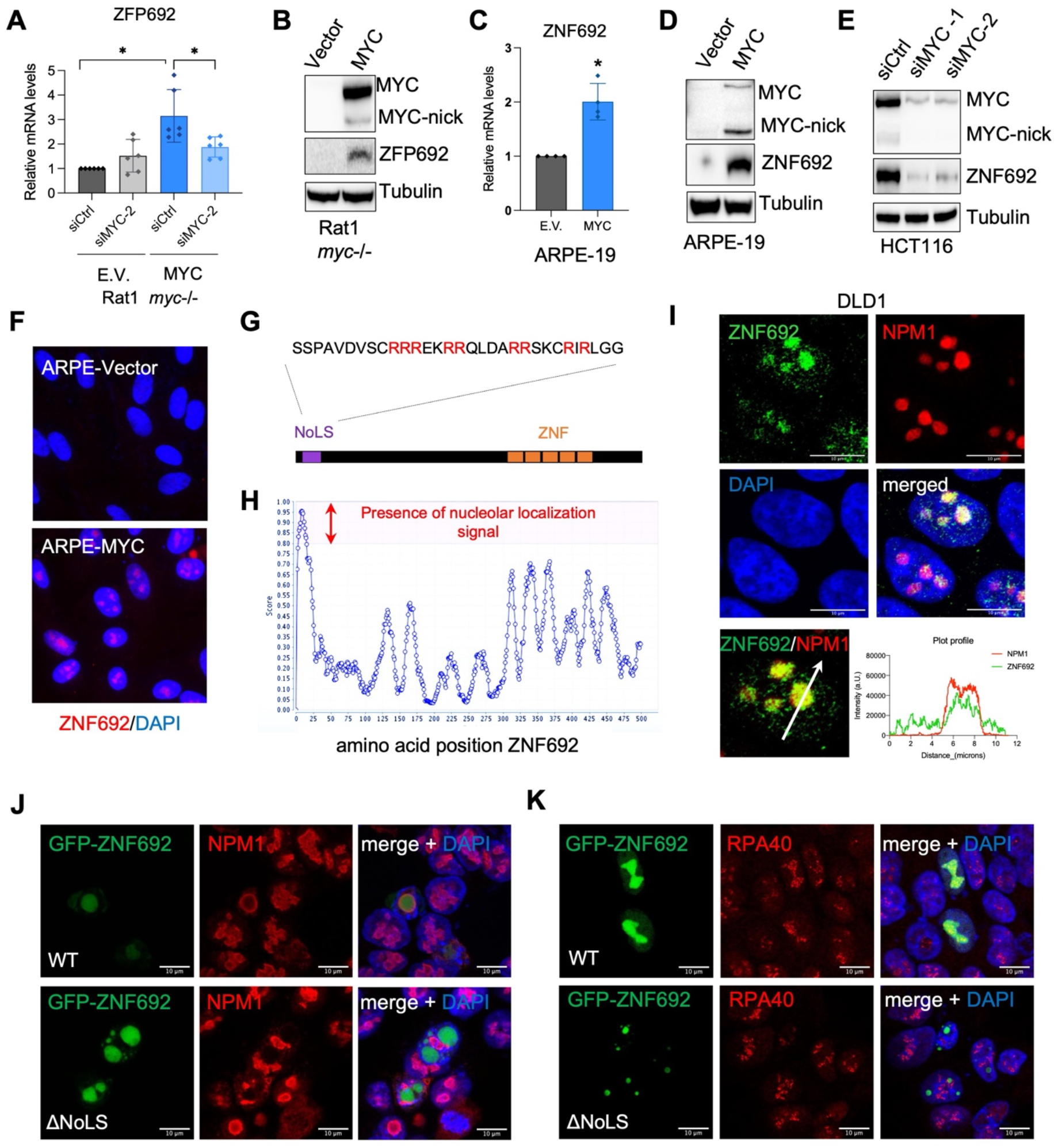
MYC promotes the transcription of ZNF692, a novel and evolutionarily conserved nucleolar factor. (A) Relative mRNA expression for ZFP692 (rodent ZNF692 orthologue) in Rat1 myc-/- cells stably expressing empty vector or MYC upon transient transfection with control siRNA (siCtrl) or siRNA for MYC. *P<0.05. (B) Western blot (WB) for ZFP692 (rodent ZNF692 orthologue) in Rat1 myc-/- cells stably expressing empty vector or MYC. (C) Relative mRNA expression for ZNF692 in ARPE cells stably expressing empty vector or MYC. *P<0.05 (D) WB for ZNF692 in ARPE cells stably expressing empty vector or MYC. (E) WB for ZNF692 in HCT116 cells transfected with control siRNA (siCtrl) or siRNA for MYC. (F) IF for ZNF692 in ARPE cells stably expressing empty vector or MYC. (G) Predicted nucleolar localization sequence (NoLS) in the N-terminus of ZNF692. (H) Identification of a high score NoLS in ZNF692 using Nucleolar Localization Sequence Detector (NoD). (I) IF showing co-localization of endogenous ZNF692 with NPM1. (J) IF of NPM1 in cells transfected with WT GFP-ZNF692 and the ΔNoLS mutant lacking the NoLS region. (K) IF of RPA40 in cells transfected with WT GFP-ZNF692 and the ΔNoLS mutant lacking the NoLS region. Deletion of the predicted NoLS relocalized ZNF692 into nucleoplasmic droplets and prevented its co-localization with NPM1 or RPA40.

ZNF692 was highly expressed in colon cancer cell lines that also had high levels of MYC in comparison with non-transformed colonic cells (HCEC) (Supplementary Fig. 1G). Knocking down *MYC* in colon cancer cells lines reduced the expression of ZNF692 (Fig. 1E and S1H). As such, colon cancer cell lines were used for further experiments to elucidate the cellular and molecular functions of ZNF692.

Immunofluorescence (IF) confirmed that MYC expression increased ZNF692 protein, which was surprisingly concentrated in the nucleoli (Fig. 1F) despite being previously proposed to function as RNA polymerase II-related transcription factor. Interestingly, using a nucleolar localization prediction tool ^37^, we identified a nucleolar localization signal (NoLS) in the N-terminal (Nt) region of ZNF692 (Fig. 1G, H). Co-IF of ZNF692 and nucleolar marker nucleophosmin 1 (NPM1) showed that ZNF692 colocalized with NPM1, confirming its localization in the nucleolus (Fig. 1I). In agreement, GFP-ZNF692 is also localized in the nucleolus (Fig. 1J) and deletion of the predicted NoLS re-localized GFP-ZNF692 into nucleoplasmic droplets that are no longer co-localized with nucleolar markers such as NPM1(Fig. 1J) and the RNAPol I component RPA40 (Fig. 1K). Sequence homology analyses demonstrated that ZNF692 emerged in chordates thus suggesting a new evolutionary acquired role (Supplementary Fig. 1I). ZNF692 contains 5 highly conserved Zn fingers domains in the C-terminal (Ct) region. The central domain of ZNF692 showed poor conservation among the analyzed species, apart from a core of 11 amino acid stretch rich in glutamic acid (E) (Supplementary Fig. 1I in red). In addition, the Nt region where the NoLS is located was also conserved in all species, thus indicating that the nucleolar localization and the potential functions of ZNF692 are evolutionary conserved.

### ZNF692 depletion reduces cell growth

Nucleolar function is highly regulated by various cellular stimuli, including nutrient availability ^38, 39^. In agreement, we found that serum stimulation, which promotes cell proliferation, caused an increase in endogenously expressed MYC and ZNF692 as well as in ectopically expressed ZNF692 (Fig. 1A) suggesting that ZNF692 can be upregulated by complementary mechanisms during growth stimulation such as transcription by MYC or increase mRNA/protein stability by growth factor stimulation. Interestingly, the increase in ZNF692 correlated with cyclin D1 (Fig. 1A) which occurs through G1-S when cells are growing and requiring more ribosomes for protein synthesis. Using siRNA-mediated knockdown (KD) of *ZNF692* we demonstrated that reducing ZNF692 levels limits the proliferation of colon cancer cells (Fig. 2B, C) in agreement with previous studies showing that ZNF692 is required for proliferation of lung, cervical, and colon cancer cell lines ^40–42^. Interestingly, knocking down *ZNF692* in MYC-expressing ARPE cells reduced their proliferation while more modest effects were observed in ARPE cells expressing empty vector (Fig. 2D, E). This is in agreement with the increased expression of ZNF692 in ARPE-MYC cells (Fig. 2D) and suggests an increased demand for ZNF692 in MYC-overexpressing cells. Prolonged *ZNF692* KD reduced proliferation in all lines tested (Supplementary Fig. 2A-B). Thus, our results demonstrated that acutely knocking down *ZNF692* caused a decrease in cell proliferation (Fig. 2B-E). Surprisingly, stable shRNA-mediated KD of *ZNF692* or stable overexpression of *ZNF692* did not significantly affect cell proliferation (Fig 2F, G) or cell size (Supplementary Fig. 2C-E) suggesting that cells adapted to the changes in expression levels of ZNF692 in cell cultures. These results led us to further investigate the importance of ZNF692 for growth *in vivo*. To ensure complete loss of function of ZNF692, we used *ZNF692* CRISPR-mediated knockout (KO) cell lines (Fig. 2H, I). Similar to stable KD of *ZNF692*, KO cells did not display a reduction in growth when assayed for a 3-day growth period *in vitro* (Fig. 2H). However, long-term growth of the same cells xenotransplanted in mice (Fig. 2J) led to the formation of smaller tumors than control cells (Fig. 2K, L and Supplementary Fig. 2F).

**Figure 2:**
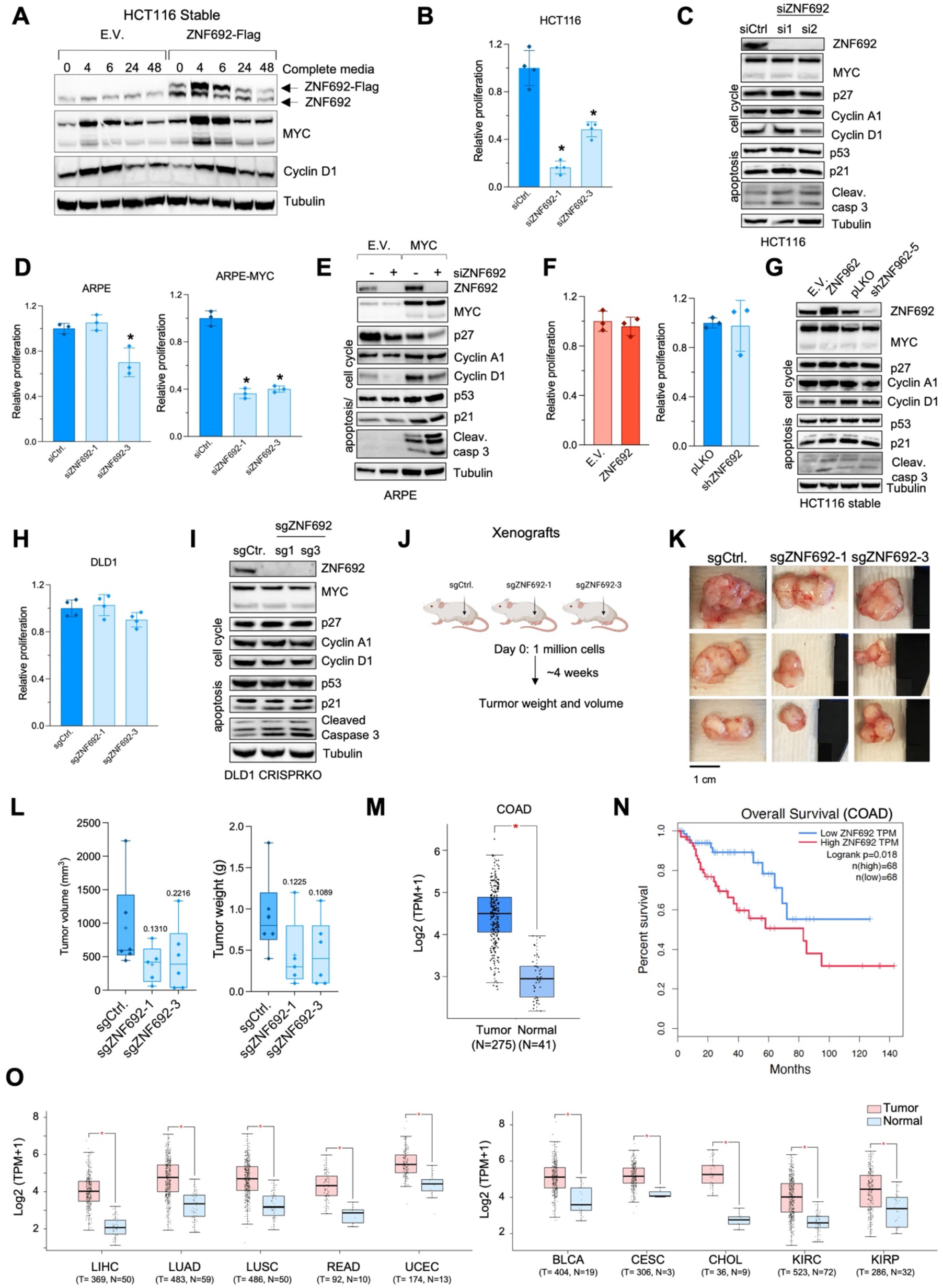
Knockdown of ZNF692 reduced the viability of proliferative cells without affecting the expression of cell cycle regulators. (A) WB of ZNF692 upregulation 4 hours after serum stimulation in parallel with MYC and Cycling D1 upregulation in stable HCT116 expressing empty vector (E.V.) or ZNF692. (B) Relative proliferation of HCT116 cells 3 days after transfection with control or ZNF692 siRNAs. *P<0.05. (C) WB for ZNF692, MYC, and markers of cell cycle and apoptosis in HCT116 cells transfected with control or ZNF692 siRNA. (D) Relative proliferation of ARPE cells stably expressing empty vector or MYC 3 days after transfection with control or ZNF692 siRNA. *P<0.05. (E) WB for ZNF692, MYC, and markers of cell cycle and apoptosis of ARPE cells stably expressing empty vector or MYC 3 days after transfection with control or ZNF692 siRNA. (F) Relative proliferation of HCT116 cells stably expressing ZNF692 (red) or shRNA for ZNF692 (blue). (G) WB for ZNF692, MYC, and markers of cell cycle and apoptosis of HCT116 cells stably expressing ZNF692 or shRNA for ZNF692. (H) Relative proliferation of DLD1 cells infected with lentiviral particles (sg1 or sg3) for ZNF692 KO or control. (I) WB for ZNF692, MYC, and markers of cell cycle and apoptosis of DLD1 cells CRISPR KO for ZNF692 or control. (J) Schematic representation of xenograft experiment with DLD1 cells CRISPR KO for ZNF692 or control. (K) Pictures of tumors collected from the xenografts experiment. (See also Supplementary figure 3F). (L) Tumor weight and volume from xenograft experiment. (M) ZNF692 mRNA expression in tumor vs normal tissues from COAD patients deposited in TCGA. (N) Patient survival correlation in relation with ZNF692 mRNA levels in COAD patients from TCGA. (O) ZNF692 mRNA expression in tumor vs normal tissues from patients of different tumor types deposited in TCGA. WB for ZNF692, MYC and markers of cell cycle and apoptosis of DLD1 xenograft tumors CRISPR KO for ZNF692 or control.

Surprisingly, as opposed to previous studies suggesting that ZNF692 regulates p27 and Cyclin D1 levels ^40, 41^, our experiments examining the expression of p27 and Cyclin D1 upon *ZNF692* siRNA-mediated KD, shRNA-mediated KD, CRISPR KO, or overexpression of *ZNF692* in multiple cell lines, found that ZNF692 had no consistent effects on p27 or Cyclin D1 levels (Fig. 2C, D, G, I, and Supplementary Fig. 2G). Moreover, KD, KO, or overexpression of *ZNF692* had no effect on the expression of other growth regulators such as p21, p53, and cyclin A1 (Fig. 2C, D, G, I, and Supplementary Fig. 2G). These results suggest that regulation of p27 and Cyclin D1 are not general functions of ZNF692, and that the ZNF692 effects on cell growth are most likely a consequence of its yet undiscovered role in the nucleolus.

By examining the expression of ZNF692 in the patient samples deposited in TCGA, we found that ZNF692 expression was dramatically elevated in colon cancer tumors when compared to normal tissues (Fig. 2M) and that elevated ZNF692 mRNA was correlated with shorter patient survival (Fig. 2N). Similar increase in ZNF692 mRNA was found in multiple other solid tumors (Fig. 2O). Taken together our results demonstrate that ZNF692 is highly expressed in tumors and is required for maximum growth of cancer cells.

### ZNF692 regulates nucleolar morphology and protein synthesis

Given that ZNF692 localized in the nucleolus, we asked whether the expression of ZNF692 would affect nucleolar morphology and activity by measuring nucleolar size and shape in relationship to ZNF692 levels. We found that tumors generated by cells in which *ZNF692* was knocked out contained smaller and rounder nucleoli (Fig. 3A, B and Supplementary Fig. 3A, B), an indication of reduced nucleolar activity and thus ribosome biogenesis. In non-perturbed cultures, we found that cells expressing elevated levels of ZNF692 (top 25%) had large nucleoli while cells expressing low levels of ZNF692 (bottom 25%) had smaller nucleoli (Fig. 3C and Supplementary Fig. 3C). Moreover, cells expressing low levels of ZNF692 displayed rounder nucleoli (Fig. 3C and Supplementary Fig. 3C). To examine the ultrastructure of the nucleolus upon *ZNF692* KD, we performed EM in cells expressing empty vector (pLKO) or shRNA for *ZNF692*. Our data showed that *ZNF692* KD caused a reduction in nucleolar perimeter as observed for xenografted cells (Fig. 3D-E and Supplementary Fig. 3D).

**Figure 3:**
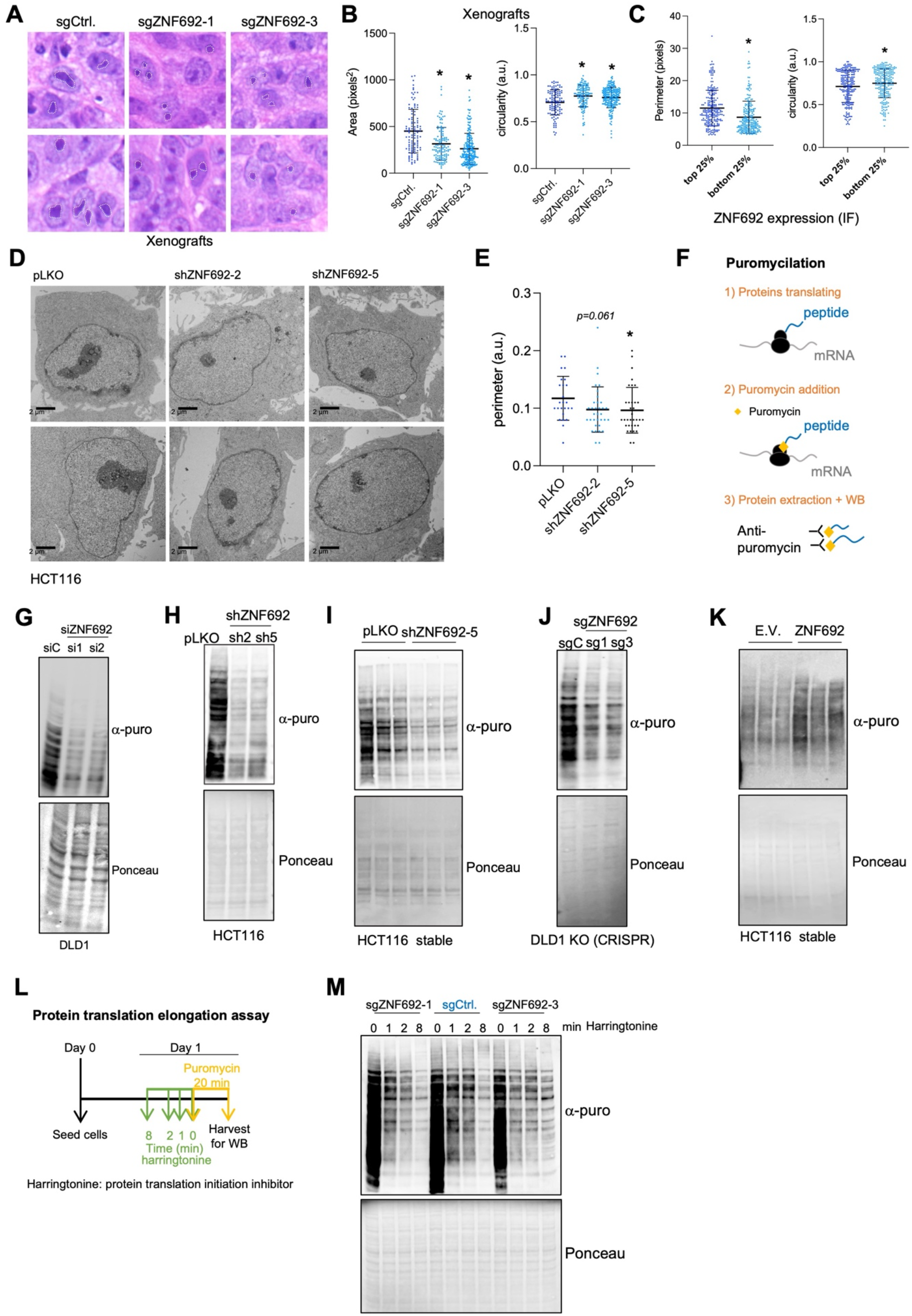
ZNF692 regulates nucleolar morphology and protein synthesis. (A) H&E staining from DLD1 xenograft tumors CRISPR KO for ZNF692 or control. White dotted line outlines the nucleolus. (See also Supplementary Fig. 3G). (B) Quantification of nucleolar area and circularity (more circular the closer to 1) of DLD1 xenograft tumors CRISPR KO for ZNF692 or control. *P<0.05. (See also Supplementary figure 3H). (C) Correlation between levels of ZNF692 measured by IF with nucleolar size (perimeter in pixels) and circularity (a.u. more circular the closer to 1) in HCT116 cells. *P<0.05. (See also Supplementary Fig. 3C) (D) Electron microscopy displaying nucleolar morphology of 2 examples of HCT116 cells after 3 days of infection with empty vector (PLKO) or shRNA for ZNF692 (#2 and #5) containing virus (See also Supplementary Fig. 2B). (E) Quantification of nucleolar area and circularity (more circular the closer to 1) from cells in C showing that ZNF692 knockdown reduces the perimeter. *P<0.05. (F) Schematic representation of the puromycilation-based assays to measure protein synthesis. (G) Puromycilation of DLD1 cells transiently transfected with control or ZNF692 siRNAs. (H) Puromycilation of HCT116 cells transiently infected with control or ZNF692 shRNAs. (I) Puromycilation of HCT116 cells stably infected with control or ZNF692 shRNAs. Three replicates are shown. (J) Puromycilation of DLD1 WT or knocked out for ZNF692 using two independent sgRNA (sg1 and sg3). These lines are a pool of cells infected with lentiviral particles containing sgRNA sg1 and sg3. (K) Puromycilation of HCT116 stably expressing ZNF692. Three replicates are shown. (L) Schematic representation of the harringtonine and puromycilation-based assay to measure translation elongation. (M) Inhibition of translation initiation with harringtonine followed by puromycilation chase in DLD1 ZNF692 CRISPR KO or control.

The exclusive nucleolar localization of ZNF692 and its correlation with nucleolar size and morphology prompted us to determine whether ZNF692 plays a role in ribosome biogenesis and/or protein synthesis. We first quantified protein synthesis upon gain and loss of function of ZNF692 using puromycilation (Fig. 3F) ^43, 44^. Knocking down *ZNF692* by siRNA (Fig. 3G, Supplementary Fig. 3E), or shRNA either transiently (Fig. 3H) or stably (Fig. 3I, Supplementary Fig. 3F) decreased protein synthesis. Moreover, knocking out *ZNF692* also reduced protein synthesis (Fig. 3J). Conversely, ectopic expression of *ZNF692* caused an increase in protein synthesis (Fig. 3K). Treating control or *ZNF692* KO cells with the inhibitor of translation initiation harringtonine prior to puromycilation (Fig. 3L), confirmed that translation elongation was reduced in *ZNF692* KO cells (Fig. 3M). ZNF692 up or down regulation did not affect the expression of nucleolar regulators such as Upstream Binding Transcription Factor (UBF), RPA40, FBL, and NPM1(Supplementary Fig. 3G-J).

### ZNF692 resides in the granular component of the nucleolus and interacts with structural and regulatory ribosome biogenesis factors

ZNF692 was proposed to function as a transcription factor due to the presence of Zn fingers in its Ct (Fig. 1G, Supplementary Fig. 1J), thus, we asked whether it regulated rDNA transcription. Using an antibody against ZNF692 we immunoprecipitated endogenous or overexpressed ZNF692 and performed chromatin immunoprecipitation (ChIP). We scanned the binding of ZNF692 to two regions on the rDNA where the transcriptional machinery binds: near the transcription start site (H1) and at the transcription end site (H13). The rDNA intergenic region (H32) where transcription machinery is not expected to bind was used as a negative control ^28^(Fig. 4A). While RNAPol I efficiently bound to rDNA at regions H1 and H13, but not to the IGS H32 region as expected, ZNF692 did not bind to any region of the rDNA regions tested (Fig 4A). Neither RNAPol I nor ZNF692 were bound to LDHA promoter which was used as an additional negative control (Fig 4A). Immunoprecipitation (IP) followed by Western blot (WB) demonstrated that the antibody for ZNF692 efficiently immunoprecipitated ZNF692 in formaldehyde-fixed lysates (Supplementary Fig. 4A). In agreement, pre-rRNA levels as measured by RT-qPCR (Fig. 4B) or IF with anti-rRNA antibody (Supplementary Fig. 4C) or RNA-FISH (Fig. S4D), were also not affected when *ZNF692* was stably knocked down (Supplementary Fig. 4C) or knocked out (Supplementary Fig. 4D). Therefore, our data showed that ZNF692 does not act as a transcription factor in the nucleolus.

**Figure 4:**
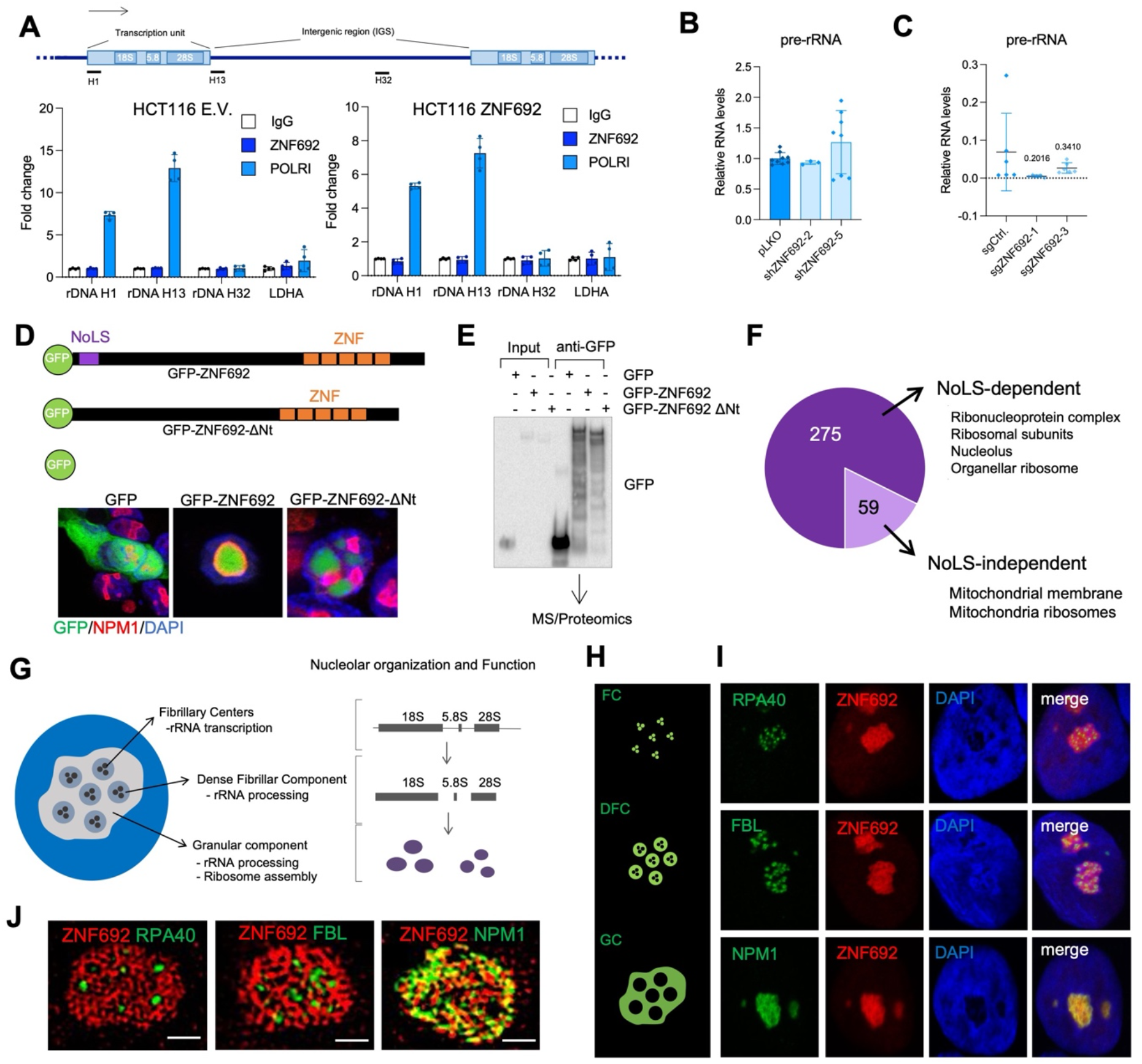
ZNF692 resides in the granular component of the nucleolus where it interacts with ribosomal proteins. (A) Chromatin immunoprecipitation (ChIP) with anti-POLRI and anti-ZNF692 and qPCR for rDNA sites H1, H13 (two different sites where the transcriptional machinery binds within the rDNA locus), H32 (intergenic regions) and LDHA promoter in HCT116 stably expressing empty vector (E.V.) or Flag-tagged ZNF692 showing that POLRI, but not ZNF692, binds to regulatory regions in the rDNA genes but not to rDNA H32 or LDHA promoter regions. Representation of two experiment with two technical replicates each out of 3 biological replicates with similar results. (B) RT-qPCR showing the pre-rRNA levels in HCT116 cells with stable ZNF692 overexpression (red) or knockdown (blue). (C) RT-qPCR of pre-rRNA in DLD1 xenograft tumors CRISPR KO for ZNF692 or control from Fig 3J-O. (D) Representation of GFP, GFP-tagged ZNF692 and ZNF692 lacking the NoLS (ΔNt) and their localization when expressed in HCT116 cells. GFP was expressed in the whole cell, WT ZNF692 colocalized with NPM1 in the nucleolus and ZNF692ΔNt localized in nucleoplasmic droplets. (E) WB for GFP comparing input and IP using anti-GFP nanotrap beads of lysates of HCT116 cells transfected with the GFP, GFP-ZNF692, and GFP-ZNF692ΔNt. Immunoprecipitants were subjected to mass spectrometry to identify ZNF692’s interactome in the nucleolus. (F) Gene ontology of ZNF692 partners that depended on the NoLS and partners that did not depend on the NoLS. (See also Supplementary Fig. 4E-F). (G) Schematic representation and description of a nucleolus and its 3 subcompartments contained inside a nucleus. (H) Representation of the expected localization of RPA40 in the fibrillary centers (FC), fibrillarin (FBL) in the dense fibrillar component (DFC) and NPM1 in the granular component (GC) of the nucleolus. (I) A representative of IF in ZNF692-transfected DLD1 cells showing the colocalization of ZNF692 with NPM1in the GC through double staining of ZNF692 (red) with RPA40, FBL, or NPM (green). (J) Structured Illumination Microscopy (SIM) in ZNF692-transfected DLD1 cells showing that ZNF692 (red) is co-localized with NPM1, but not RPA40 and FBL (green). (See also Supplementary Fig. 4E).

To investigate the molecular function of ZNF692 in the nucleolus, we identified ZNF692’s interactome. Lysates of cells transfected with either GFP, GFP-ZNF692 WT, or GFP-ZNF692 ΔNt (lacking the NoLS) (Fig. 4D) were immunoprecipitated with GFP-nanotrap beads and the co-IP proteins were subjected to mass spectrometry (Fig. 4E). To identify nucleolar-specific ZNF692 interactors, only proteins that lost their binding to ZNF692 when the NoLS was deleted were considered (see methods). This approach identified 334 proteins as ZNF692 interactors; with the majority of these interactions (275) dependent on the NoLS. Consistently with the nucleolar localization of ZNF692, gene ontology analysis determined that its interactors are ribosomal proteins and rRNA processing factors (Fig. 4F, Supplementary Fig. 4E, F).

By performing co-IF of ZNF692 with RPA40 (a marker of the FC), Fibrillarin (FBL, a marker of the DFC) and NPM1 (a marker of the GC) (Fig 4G-H), we found that ZNF692 was excluded from the FC and the DFC, and predominantly (but not completely) co-localized with NPM1 in the GC (Fig. 4I). Structure illumination microscopy (SIM) found that ZNF692 consistently separated from RPA40 and FBL while mostly co-localized with NPM1 (Fig. 4J and Supplementary Fig. 4I) confirming that the GC is the primary sub-nucleolar residence of ZNF692. The absence of ZNF692 from the FC is in agreement with its inability to bind rDNA and to regulate its transcription. Thus, the localization of ZNF692 in the GC of the nucleolus suggests that ZNF692 is involved in ribosome subunit maturation (Fig. 4G).

### ZNF692 interacts with components of the small subunit processome and the exosome complex

ZNF692 is bound to a group of 16 regulatory proteins involved in rRNA processing and ribosome assembly (Fig. 5A and Supplementary Table 1). Eleven of these interactors are involved in 18S rRNA maturation and 4 of them in the 28S rRNA maturation. Several of the ZNF692 interacting factors were found to be components the 90S small subunit processome in yeast ^45^, including the 18S rRNA processing factors KRR1, the KRR1 interactor RCL1, IMP3, and DIMT1.

**Figure 5:**
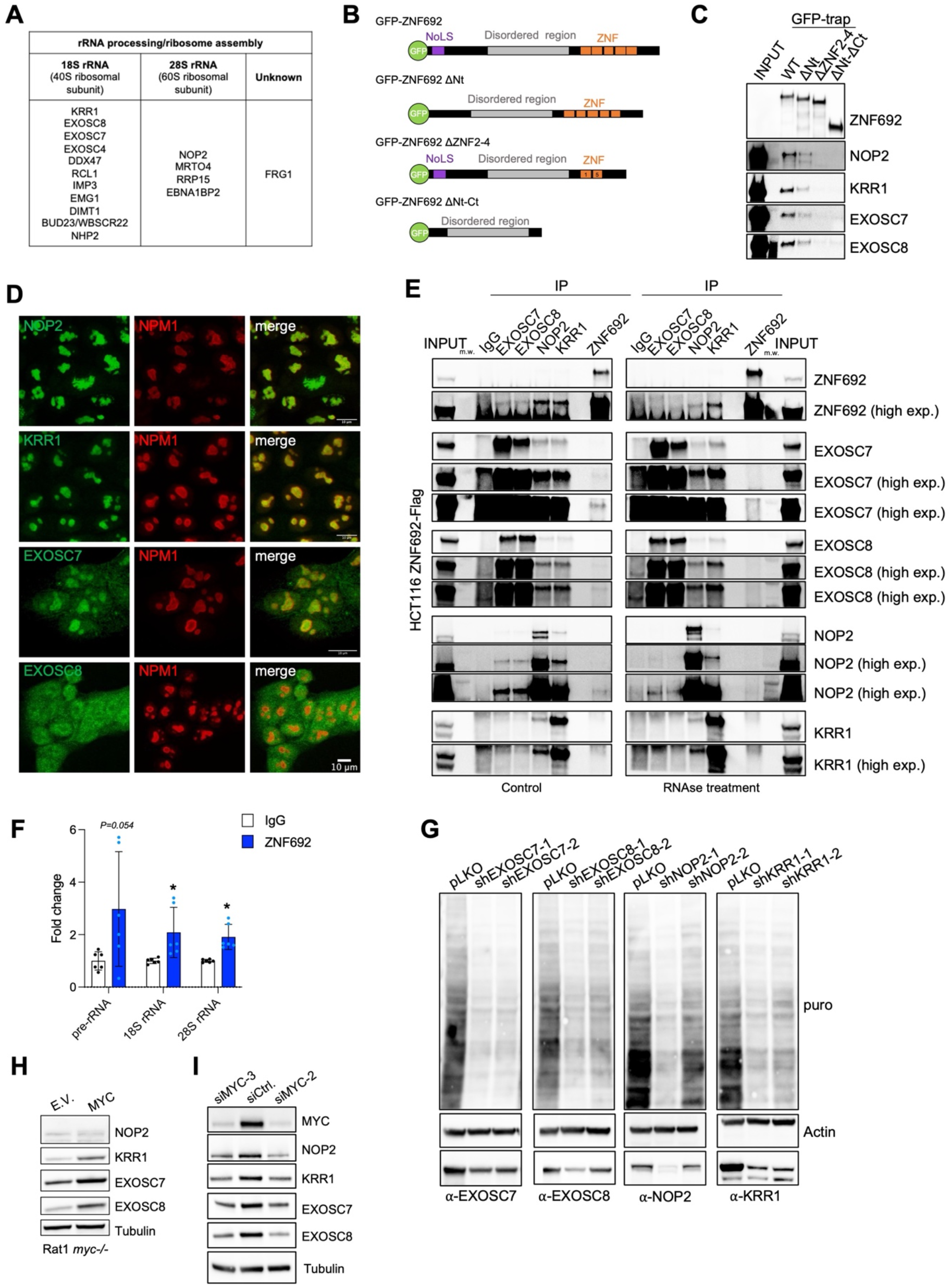
ZNF692 interacts with the components of the exosome complex. (A) ZNF692 interactors and their function in ribosomal subunit assembly. (See also Supplementary Table 1). (B) Schematic representation of ZNF692 recombinant proteins purified from insect cells to be used for *in vitro* assays. (C) IP of recombinant ZNF692 using GFP nanotrap beads in the presence of nuclear extracts of DLD1 cells followed by immunoblot for NOP2, KRR1, EXOSC7, and EXOSC8. (D) IF of NOP2, KRR1, EXOSC7, or EXOSC8 together with NPM1 showing their colocalization with NPM1 in the granular component of the nucleolus. (E) IP of the endogenous EXOSC7, EXOSC8 NOP2, KRR1, and ZNF692 of HCT116 cells stably overexpressing ZNF692 followed by immunoblot for ZNF692, NOP2, KRR1, EXOSC7, and EXOSC8. Left panel, control; right panel lysates were treated with 20 µg/ml RNase A prior to IP. (F) RNA**-**IP of ZNF692 in DLD1 cells on rRNA, ZNF692 binds rRNA. Three independent IPs with two technical replicates each are represented. (G) Puromycilation in DLD1 cells 3 days after infection with lentiviral particles containing shRNA for either control or NOP2, KRR1, EXOSC7, or EXOSC8. (H) WB for NOP2, KRR1, EXOSC7 and EXOSC8 in cells *myc-/-* fibroblasts expressing empty vector (E.V.) or MYC. (I) WB for MYC, NOP2, KRR1, EXOSC7, and EXOSC8 in lysates of DLD1 cells transiently transfected with siRNA for MYC or control.

To validate the interactions identified by proteomics, we performed Co-IPs of ZNF692 with the exosome complex components EXOSC7 and EXOSC8; the rRNA methyltransferases NOP2 ^46^, BUD23 ^47^ and DIMT1 ^48^; the processome factor KRR1; and the pseudouridylase NHP2 ^49^. Using GFP nanotrap beads and purified GFP-ZNF692 WT or mutants lacking the Nt including the NoLS, lacking the 2-4 Zn fingers in the Ct and lacking both the Nt and Ct regions (Fig. 5B), we co-IP ZNF692 binding partners from nuclear lysates. In this assay, ZNF692 was bound to NOP2, KRR1, EXOSC7, and EXOSC8 (Fig. 5C) but not to NHP2, BUD23, or DIMT1 (Supplementary Fig. 5A). Deletion of the Nt domain of ZNF692 reduced its binding to NOP2, KRR1, EXOSC7, and EXOSC8 while disruption of the Ct Zn fingers abolished it. In agreement with the co-IP results, co-IF with NPM1 demonstrated that NOP2, KRR1, and EXOSC7 localize in the GC of the nucleolus as seen by their colocalization with NPM1 where ZNF692 resides (Fig. 5D). EXOSC8 was present in the cytoplasm and nucleus (Fig. 5D). NHP2 did not localize in the GC of the nucleolus (Fig S5B), thus validating its lack of co-IP with ZNF692. Reciprocal co-IPs for endogenous EXOSC7, EXOSC8, NOP2, KRR1 with ZNF692 in ZNF692-overexpressing cells confirmed that ZNF692 interacts with EXOSC7, EXOSC8, NOP2, and KRR1 (Fig. 5E). Interestingly, our results showed that EXOSC7, EXOSC8, NOP2, and KRR1 also interacted with each other (Fig. 5E), thus suggesting the formation of a larger complex containing the exosome and components of the small subunit processome in mammalian cells which had not been previously described.

Given that deletion of the Zn fingers reduced the binding of ZNF692 with its interactors (Fig. 5C), we asked whether the presence of rRNA could be involved in forming a complex containing ZNF692 and its rRNA processing interactors. Indeed, the interaction between ZNF692, EXOSC7, EXOSC8, and NOP2 depended on the presence of RNA because treating the lysates with RNAse A prevented their reciprocal interaction as measured by IP (Fig. 5E). Conversely, the interaction between ZNF692 and KRR1 remained unaffected even when RNA was degraded (Fig. 5E). RNA-IP with an anti-ZNF692 using UV-crosslinked cells overexpressing ZNF692 demonstrated that ZNF692 interacted with the 18S and 28S rRNA (Fig. 5F). NPM1, which is known to bind rRNA was used as a positive control (Supplementary Fig. 5C). Furthermore, treatment with 0.05 µg/ml of Actinomycin D, known to specifically inhibit rDNA transcription (Supplementary Fig. 5D, E), led to the relocalization of ZNF692 from the nucleolus to the nucleoplasm (Supplementary Fig. 5E), suggesting that the presence of rRNA contributes to the retention of ZNF692 in the nucleolus.

To determine whether ZNF692 interactors affected protein synthesis as a consequence of their expected role on ribosome biogenesis ^46, 50, 51^, we measured translation by puromycilation upon transient KD of *EXOSC7*, *EXOSC8*, *NOP2*, and *KRR1*. KD of all 4 genes led to a reduction in protein synthesis as measured by puromycilation (Fig. 5G). Similar to ZNF692, stably knocking down *EXOSC7*, *EXOSC8*, *NOP2*, and *KRR1* had variable and often modest effects on cell proliferation, indicating that reduction in protein synthesis caused by downregulation of these genes is not a consequence of cell death (Supplementary Fig. 5F).

ZNF692 interacting partners KRR1, EXOSC7, and EXOSC8 similar to ZNF692, were also upregulated in MYC-expressing fibroblasts (Fig. 5H). Knocking down *MYC* in colon cancer cells decreased the expression of NOP2, KRR1, EXOSC7 and EXOSC8 (Fig. 5I). These results suggest that the members of the complex containing ZNF692 and ribosome maturation factors are co-upregulated by MYC in rapidly proliferating cells. Moreover, the expression of KRR1, EXOSC7, EXOSC8, and NOP2 was also upregulated in colon cancer samples deposited in the TCGA (Supplementary Fig. 5G). ZNF692 was the most upregulated gene in this group when compared to the basal levels of each gene in normal tissues. Interestingly, while high ZNF692 mRNA expression levels correlated with poor COAD patient survival (Fig. 2N), the expression of KRR1, EXOSC7, EXOSC8, and NOP2 did not correlate with COAD patient survival (Supplementary Fig. 5H). These results suggest that ZNF692 is a nucleolar adaptor that facilitates ribosome biogenesis in hyperproliferative mammalian cells.

### ZNF692 promotes 18S maturation and the formation of functional small ribosomal subunit

Three out of the four mature rRNAs are transcribed as a single polycistronic pre-rRNA (47S), which is processed to generate 18S, 5.8S, and 28S (Fig. 6A). The molecular details guiding the final steps of the 18S rRNA maturation in mammalian cells are not yet fully mapped. Given that ZNF692 interacted with rRNA processing factors, we investigated whether altering ZNF692 levels would affect the processing of rRNA. For that, we performed Northern blot of total RNA extracted from control or cells in which *ZNF692* was KD or KO using the rRNA probes 5’-ETS, mapping the pre-rRNA in the initial 5’ region, ITS1 mapping the internal transcribed spacers between 18S and 5.8S rRNA regions, and ITS2 mapping the internal transcribed spacers between 5.8S and 28S rRNA regions (Fig. 6A) ^50^. The results showed that *ZNF692* depletion led to an accumulation of the 21S, the precursors of 18S rRNA, visualized by the ITS1 probe (Fig. 6B, C. and Supplementary Fig. 6A, B, E). In contrast, there were no alterations in the rRNA processing steps mapped with the 5’-ETS and ITS2 probes (Supplementary Fig. 6C, D).

**Figure 6:**
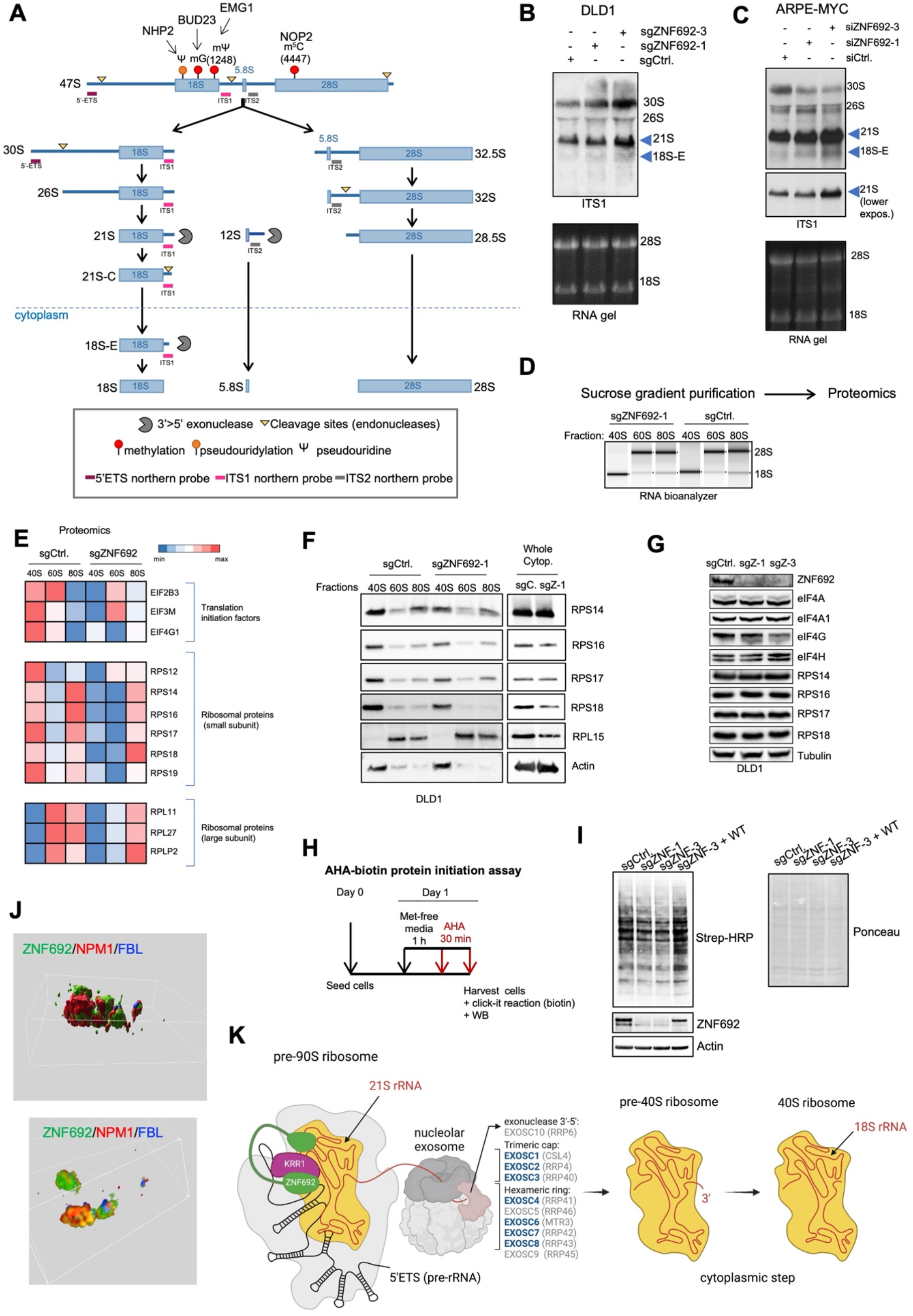
ZNF692 enhances 18S rRNA processing and small ribosomal subunit assembly. (A) Schematic representation of the pre-rRNA processing pathway showing the functions of ZNF692 interactors. (B) Northern blot of control of ZNF692 CRIPSR KO DLD1 cells. Blue arrows indicate the 21S and 18S-E. (C) Northern blot of ARPE cells stably expressing MYC and transiently transfected with siRNA for ZNF692 or control. Blue arrows indicate the 21S and 18S-E fragments. (D) Schematic representation of the proteomics experiments from 40S, 60S, and 80S cytoplasmic fractions collected of DLD1 KO for ZNF692 or control. (E) Proteomics analysis heatmap showing fold change between DLD1 ZNF692 KO and control from 40S, 60S, and 80S single ribosomes collected from cytoplasmic fractions. (F) WB of cytoplasmic 40S, 60S, and 80S fractions of DLD1 ZNF692 KO or control. (G) WB of total cell lysates ZNF692KO or control for proteins identified in (E) and (F). Total expression levels of these factors are not altered by ZNF692 KO. (H) Schematic representation of Click-IT^TM^ AHA experiments. (I) Click-IT^TM^ AHA experiments in DLD1 *ZNF692* KO cells (-/+ ectopic ZNF692) or control cells. (J) 3D reconstruction of immunofluorescence images (Fig. S6F) for NPM1, ZNF692, and FBL in DLD1 KO cells stably overexpressing ZNF692 showing the surface (upper panel) and the volume (lower panel) of ZNF692, NPM1, and FBL in the nucleolus. (See also Supplementary Fig. 6G and 6H). (K) Model of the interaction of ZNF692 with KRR1 and rRNA in the 90S processome complex together with the exosome complex.

Components of the processome such as RCL1 are necessary for the production of 21S, 21S-C and 18S-E intermediates ^23^. Krr1 was found to be necessary for the formation of the small subunit and the processing of the 18S rRNA in yeast ^51–53^. Moreover, *Krr1* is elevated in mouse pluripotent stem cells and its KD decreases small ribosomal subunit assembly ^54^, thus linking Krr1 to increased protein synthesis and proliferation. Given that ZNF692 directly interacts with factors of the small subunit processome and recruits the exosome complex (Fig. 5E), the accumulation of the 21S and 18S-E rRNA upon *ZNF692* KO or KD suggested that ZNF692 is involved in 18S pre-rRNA processing, defining a mechanism for the decrease in protein synthesis upon *ZNF692* KD or KO (Fig. 3G-M).

To directly test whether the biogenesis of the ribosomal subunit was affected upon *ZNF692* KO, we performed proteomics of ribosomes subunits 40S (small), 60S (large) and whole ribosomes 80S from the cytoplasm (Fig. 6D). Our results demonstrated that the presence of translation initiation factors, and ribosomal proteins were decreased when *ZNF692* was knocked out predominantly in the 40S ribosomes (Fig. 6E) which is in agreement with ZNF692’s role in the processing of the 18S rRNA. We validated by WB that RPS14, RPS16, RPS17, and RPS18 were decreased in the 40S cytoplasmic fractions of *ZNF692* KO cells in comparison with control cells (Fig. 6F). Importantly, the expression of these proteins was not altered by ZNF692 KO (Fig. 6G). RPL15 was only present on the 60S and 80S as expected and its abundance did not change in *ZNF692* KO cells (Fig. 6F-G). Actin was used as loading control. Krr1 and Rps14 interaction has been found to be important for small ribosomal subunit maturation in yeast ^52^. Our results indicate that ZNF692 by interacting with KRR1 plays a role in this KRR1-RPS14 interaction and the final maturation steps of the small ribosomal subunit. Delayed maturation of the small ribosomal subunit may lead to defects in translation initiation and thus defects in overall protein synthesis.

Translation initiation in eukaryotes involves the formation of a 43S pre-initiation complex containing the 40S mature ribosomal subunit and multiple translation initiation factors including eIF1, eIF1A, eIF3, eIF5, and the ternary complex eIF2-GTP-Met-tRNA ^55^. Once the 43S pre-initiation complex is formed, mRNA is recruited by the eIF4F complex (containing eIF4E, eIF4A, eIF4G, eIF4A). Then, the 40S scans the mRNA to find the translation initiation codon at which point the 60S ribosomal subunit is recruited to start protein elongation ^56^. Interestingly, structural ribosomal proteins such as Rps16 play regulatory roles in translation ^57^. The reduction of eIF2B3, eIF4G1, and eIF3M and RPS16 and other RPS proteins in the 40S fraction of *ZNF692* KO cells (Fig. 6E) suggest that the formation of the 43S pre-initiation complex is more efficient in cells containing ZNF692. To test whether the absence of ZNF692 affected translation initiation, we performed Click-iT^TM^ AHA incorporation experiments to measure nascent protein synthesis in *ZNF692* KO cells or control cells (Fig. 6H). Our results showed that the absence of ZNF692 reduced AHA incorporation and that reconstitution of ZNF692 rescued it (Fig. 6I and Supplementary Fig. 6F.) Thus, our data suggests that ZNF692 by facilitating the maturation of the small ribosomal subunit, enhances translation initiation and thus protein synthesis in highly proliferative cells.

Using SIM super resolution microscopy for ZNF692 and the universal nucleolar markers NPM1 and FBL, we found that ZNF692 partially co-localized with NPM1 (Fig. 6J and Supplementary Fig. 6G, H) and that there were regions in the GC where ZNF692 was enriched forming sub-nucleolar domains within the GC (Fig. 6J and Supplementary Fig. 6G, H). These results support the model that ZNF692 creates a hub in the GC of the nucleolus specialized in the maturation and assembly of the small ribosomal subunit before its export to the cytoplasm. By analyzing ZNF692 structure ^58^, we found that ZNF692 has a central disordered region that may facilitate the proximity between the Nt and Ct (Supplementary Fig. 6I). The presence of a central disordered region was confirmed by Protein DisOrder prediction System (PrDOS) (Supplementary Fig. 6J). It is possible that the central disorder region in ZNF692 (Supplementary Fig. 6J) participates in the phase separation properties of the nucleolus allowing it to segregate into a specialized compartment. In this sub-compartment, ZNF692 bridges pre-rRNA to KRR1 and the exosome complex and thus, enhances the maturation of 21S rRNA to 18S rRNA and thus the small ribosome subunit (Fig. 6K).

## Discussion

rRNA processing and ribosome biogenesis have been extensively studied in yeast. However, specialized molecules that regulate the more complex nucleolar functions in mammalian cells are not well understood. ZNF692 evolved in chordates and is therefore absent in yeast. Interestingly ZNF692 is key to physically and functionally connect conserved multiprotein complexes necessary for ribosome biogenesis. Thus, providing information on specific steps and players specialized for ribosome biogenesis in mammalian cells.

Given that highly proliferative cells have high demands for protein synthesis to grow and divide, it is not surprising that ribosome biogenesis and tumorigenesis are intimately linked. MYC has been causally related to tumor initiation and maintenance in part due to its potent capacity to boost ribosome biogenesis ^26^. MYC upregulates genes involved in all of the steps of ribosome biogenesis from rDNA transcription to rRNA maturation and genes encoding ribosomal proteins ^9, 59^. This connection between ribosome biogenesis and tumorigenesis is also apparent in genetic disorders with predisposition to tumor development. For example, Diamond-Blackfan anemia (DBA), a dominant autosomal bone marrow failure syndrome, has been correlated with several ribosomal protein gene mutations. Patients with DBA have a significantly higher risk of developing cancers of various types, including AML, colon cancer, and osteosarcoma ^60, 61^. Interestingly, mutations in ribosomal proteins such as RPS19, RPL11, RPL5, or RPL35a are linked to the development of cancer ^62^. Because ribosomal proteins and MYC are essential for normal cell activity, targeting these proteins with the goal of treating cancer could have detrimental effects to normal cells and tissues. Our study demonstrates that ZNF692 is a new nucleolar regulator, which is highly induced by MYC and enhances protein synthesis. We found that that ZNF692 increases translation efficiency of in highly proliferating cells, possibly becoming an Achilles’ heel in tumors

Microscopy shows that ZNF692 forms a subdomain in the GC that is likely important for the final steps of rRNA maturation and ribosome biogenesis. Indeed, proteomics analysis indicates that ZNF692 interacts with ribosomal protein and factors involved in the maturation of small ribosome subunit such as KRR1 and the components of the exosome complex. In agreement with the formation of a complex containing ZNF692 and the exosome complex, cells deficient in ZNF692 present an accumulation of the 21S rRNA. These results suggest that acts as a scaffold facilitating the formation of a macrocomplex containing the processome, the exosome, and pre-rRNA to enhance 18S rRNA and small ribosome subunit maturation.

In summary, our uncovers a functionally specialized sub-compartment in the granular component of the nucleolus responsible for small ribosome subunit maturation as a novel step in translation regulation used by cancer cells to cells align translation efficiency with the increased transcription promoted by MYC. While most nucleolar proteins have constitutive expression and activity, our data show that ZNF692 senses and responds to growth signals to fine tune nucleolar activity with the growth state of cells, thus making ZNF692 an interesting entry point to better understand nucleolar activity in hyperproliferative mammalian cells.

## ACKNOWLEGEMENT

We are grateful to the Sorrell lab members, Drs. Phillip Scherer and Jonathan Friedman for their feedback. This research was supported by Cancer Prevention and Research Institute of Texas (CPRIT) RR170063 to JBW and RR150059 to MS, NIH (R01GM125812) to MB, American Cancer Society IRG-17-174-13, Welch I-1914, NCI R01CA245548, UTSW Kidney Cancer SPORE Career Enhancement Program [P50CA196516/MCS] and NIH (R35CA197311 and the Welch Foundation (I-1961 to JTM. MCS is the Virginia Murchison Linthicum Scholar in Medical Research. Training fellowship supported by a Hamon Center for Regenerative Science and Medicine to MLN. IB is supported by training grant RPpqrrsq. The authors acknowledge the UT Southwestern Live Cell Imaging Facility (Harold C. Simmons Cancer Center) supported in part by an NCI Cancer Center Support Grant, 1P30 CA142543-01.

## AUTHOR CONTRIBUTIONS

CJ, MLN, YHH and MCS planned the experiments and wrote the manuscript; CJ, MLN, and YH performed most of the experiments, WS and JBW purified recombinant proteins and performed the in vitro phase separation assay, JTM and T-CC performed Northern blots. DM assisted with SIM imaging, SB with bioinformatics, YHH, MB and JTM provided reagents, protocols and input.

## Supplementary figures

**Supplementary Figure 1:**
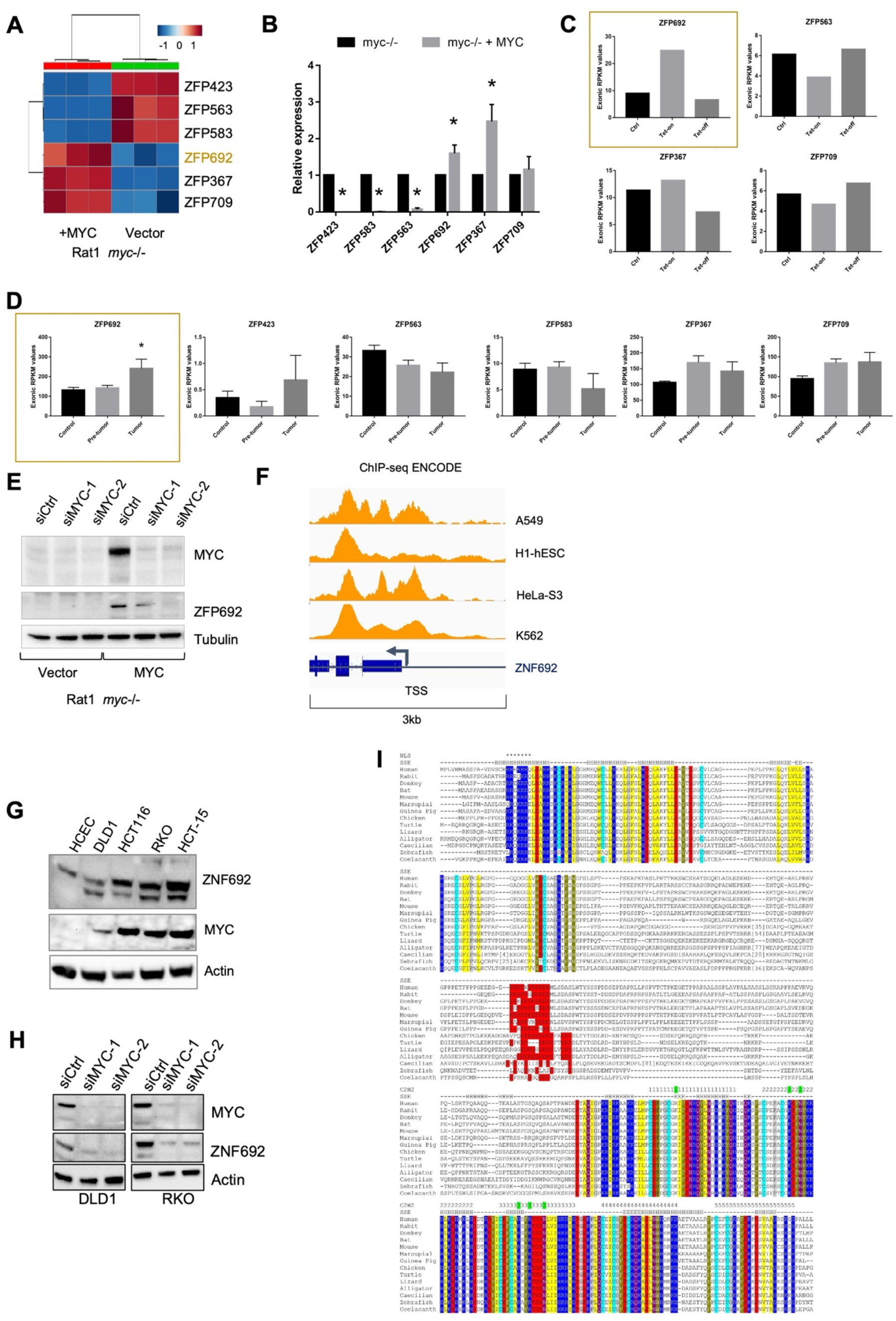
MYC regulates the expression of zinc finger-containing proteins (ZNF). (A) Heatmap of Zn finger-containing genes identified through RNA-seq comparing Rat1 *myc-/-* fibroblasts expressing empty vector or reconstituted with human MYC (hMYC). (B) Relative mRNA for all Zn fingers proteins in (A) in Rat1 *myc-/-* fibroblasts expressing empty vector or reconstituted with human MYC (hMYC). (C) Relative mRNA of Zn finger-containing genes found in (A) from previous publication ^35^ comparing WT and a mouse model of liver carcinoma driven by an inducible Tet-MYC transgene. MYC is overexpressed by tetracycline (Tet-on) but not in the absence of tetracycline (Tet-off). Highlighted is ZFP692, ZNF692 orthologue in rodents. (D) Relative mRNA ZNF692 expression in control, pre-tumoral and tumor samples from Eµ-MYC mice. Data extracted from previous publication ^36^. Highlighted is ZFP692, ZNF692 orthologue in rodents. (E) WB for ZFP692 in Rat1 *myc^−/−^* fibroblast expressing vector or hMYC 3 days after transfection with control or MYC siRNA. (F) MYC peaks from ENCODE ChIP-seq data (processed with the hg19/reference genome) on the regulatory regions of ZNF692 in different cell lines. (G) WBs comparing the expression levels of ZNF692 in non-transformed colonic HCEC and several colon cancer cell lines. (H) WB for ZNF692 in DLD1 and RKO colon cancer cells 3 days after transfection with control or MYC siRNA. (I) Multi-species comparison of ZNF692 protein showing domain and amino acid conservation. ZNF692 first appeared in chordates.

**Supplementary Figure 2:**
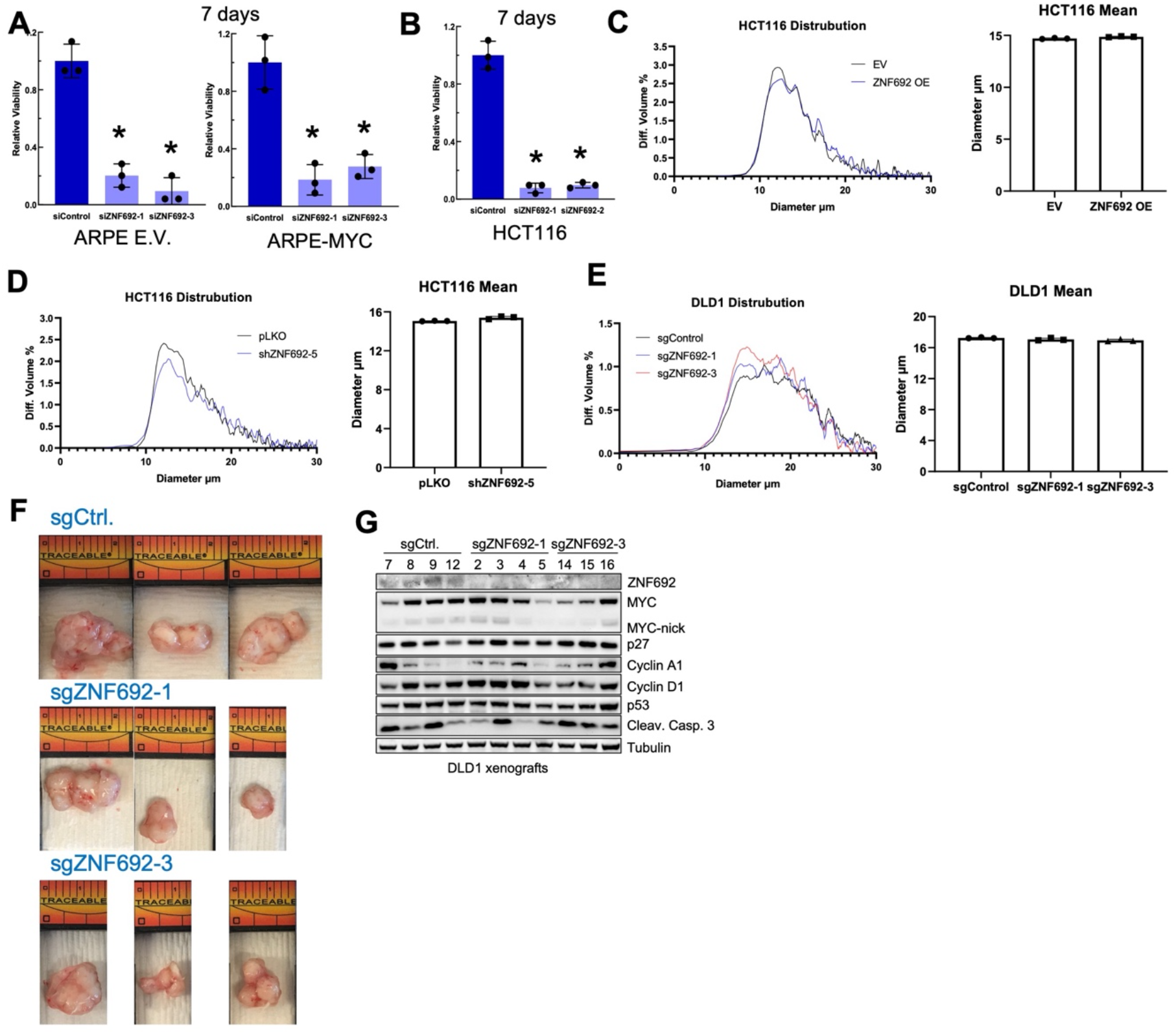
Transient knockdown of ZNF692 reduced the viability of proliferative cells without affecting the expression of cell cycle regulators. (A) Relative proliferation of ARPE cells expressing empty vector or MYC 6 days after transfection with control or ZNF692 siRNA. (B) Relative proliferation of HCT116 cells 6 days after transfection with control siRNA or ZNF692 siRNA. (C) Cell size distribution measured by Beckman Coulter Z2 Particle Count and Size Analyzer in HCT116 cell expressing empty vector (EV) or ZNF692. (D) Cell size distribution measured by Beckman Coulter Z2 Particle Count and Size Analyzer in HCT116 expressing empty vector (pLKO) or ZNF692 shRNA. (E) Cell size distribution measured by Beckman Coulter Z2 Particle Count and Size Analyzer in DLD1 control or ZNF692 CRIPSR KO cells. (F) Pictures from DLD1 tumor xenografts ZNF692 KO or control. (G) Quantification of the DLD1 tumor xenografts ZNF692 KO or control nucleoli perimeter from H&E staining.

**Supplementary Figure 3:**
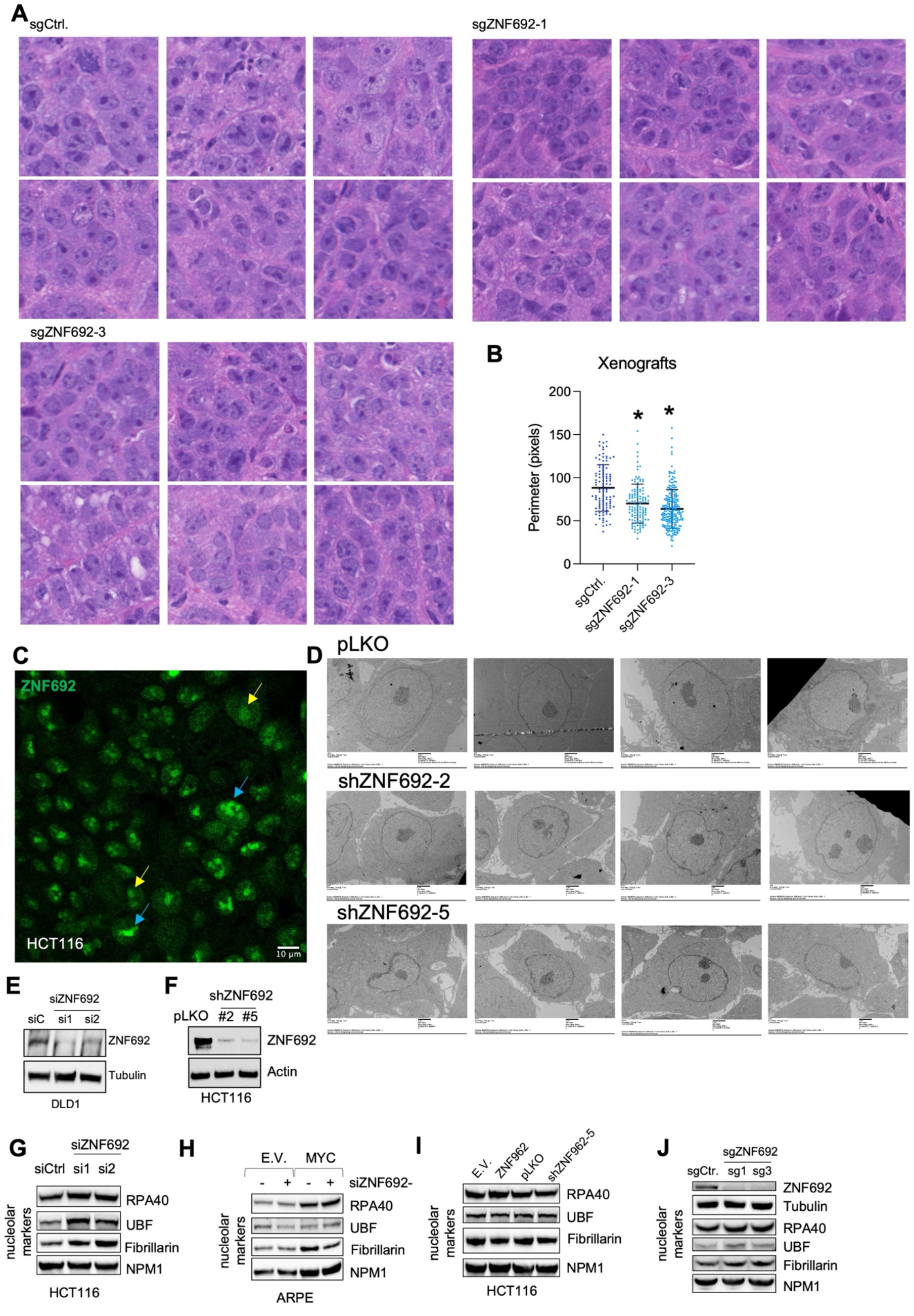
ZNF692 regulates nucleolar morphology and protein synthesis. (A) H&E staining of DLD1 tumor xenografts ZNF692 KO or control. (B) Quantification of nucleolar perimeter of DLD1 xenograft tumors CRISPR KO for ZNF692 or control. *P<0.05. (C) IF of ZNF692 in HCT116. Yellow arrows point to low ZNF692 expression cells, blue arrows point to high ZNF692 expression cells. (D) EM images of HCT116 cells expressing empty vectors or shRNA for ZNF692. (E) WB validating of ZNF692 knockdown in DLD1 transfected with siRNA. (F) WB validating downregulation of ZNF692 by shRNA in stable HCT116 cells. (G) WB of HCT116 cells 3 days after transfection with control or ZNF692 siRNAs. (H) WB of ARPE expressing empty vector (E.V.) or MYC cells transiently transfected with control or ZNF692 siRNAs. (I) WB of HCT116 cells stably expressing empty vector (E.V.) or ZNF692 or control (pLKO) or ZNF692 shRNA. (J) WB of control or ZNF692 CRIPSR KO DLD1 cells.

**Supplementary Figure 4:**
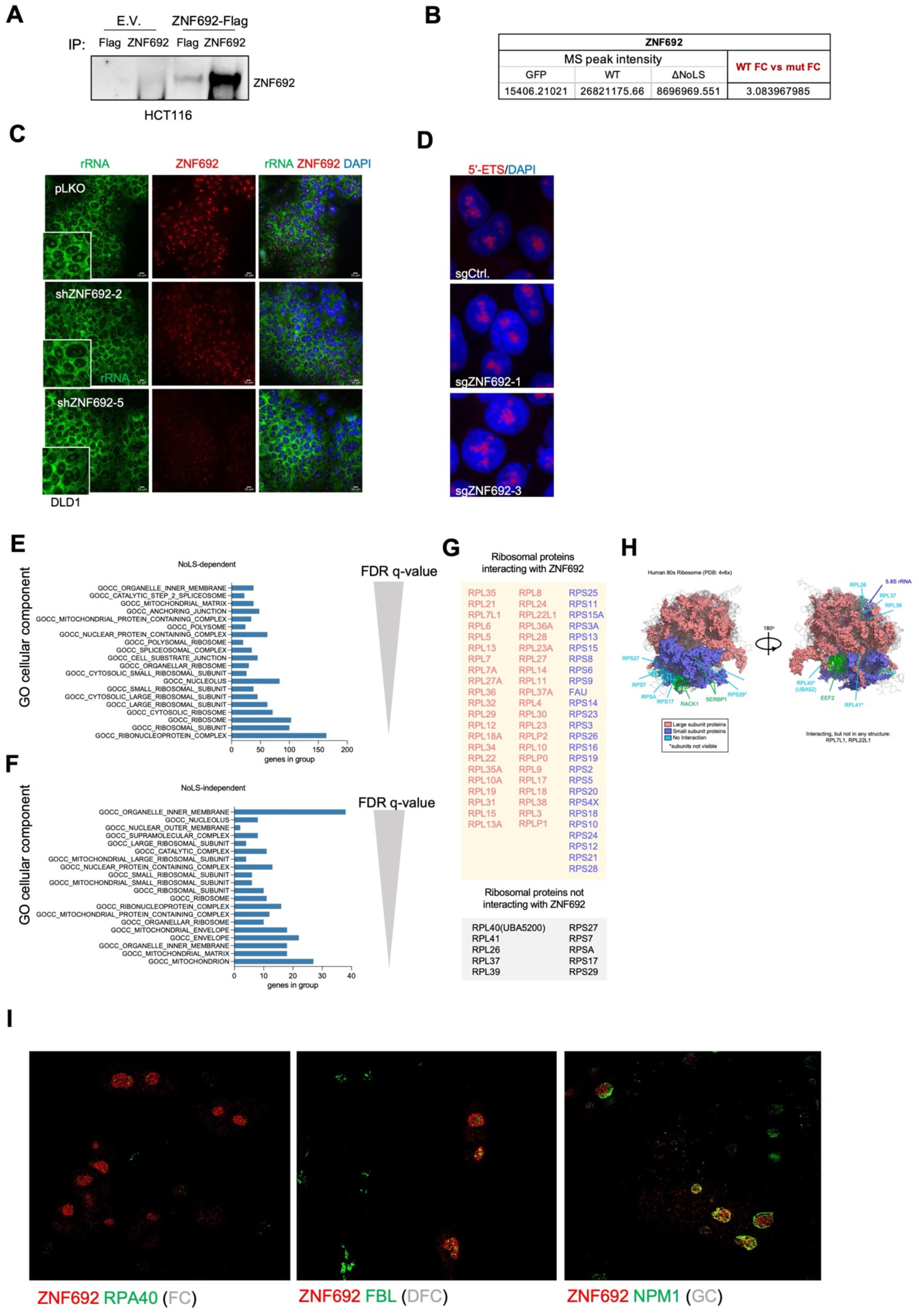
ZNF692 resides in the granular component of the nucleolus where it interacts with ribosomal proteins. (A) Immunoprecipitation of formaldehyde-fixed lysates from HCT116 stably expressing empty vector (E.V.) or Flag-tagged ZNF692 using anti-Flag and anti-ZNF692 and immunoblot with anti-ZNF692 showing that anti-ZNF692 immunoprecipitated ZNF692 in fixed lysates. (B) ZNF692 peak intensity identified in the mass spectrometry pull-down shown in Fig. 4. WT ZNF692 was present at ∼3.1 fold higher than the ZNF692 ΔNoLS. (C) IF using anti-rRNA and ZNF692 antibody showing equal rRNA presence in the nucleolus and in the cytoplasm of in HCT116 cells with stable shRNA for ZNF692 or control. (D) RNA FISH for 5-ETS (targeting pre-rRNA) in DLD1 cells CRISPR KO for ZNF692 or control. (E) Gene ontology analysis of the ZNF692 interactors that depend on the NoLS of ZNF692 ranked by FDR-value. (F) Gene ontology analysis of the ZNF692 interactors that do not depend on the NoLS of ZNF692 ranked by FDR-value. (G) Table containing ribosomal proteins found by mass spectrometry to interact with ZNF692. Components of the large subunits in red and components of the small subunit in blue. Ribosomal proteins known to be part of ribosomes that were not found to interact with ZNF692 are listed in black. (H) Structural reconstruction of the mature ribosome containing the ribosomal proteins found to interact with ZNF692. (I) Structured Illumination Microscopy (SIM) of DLD1 showing that ZNF692 is co-localized with NPM1, but not RPA40 and FIB, low magnification of the image shown in Figure 4J.

**Supplementary Figure 5:**
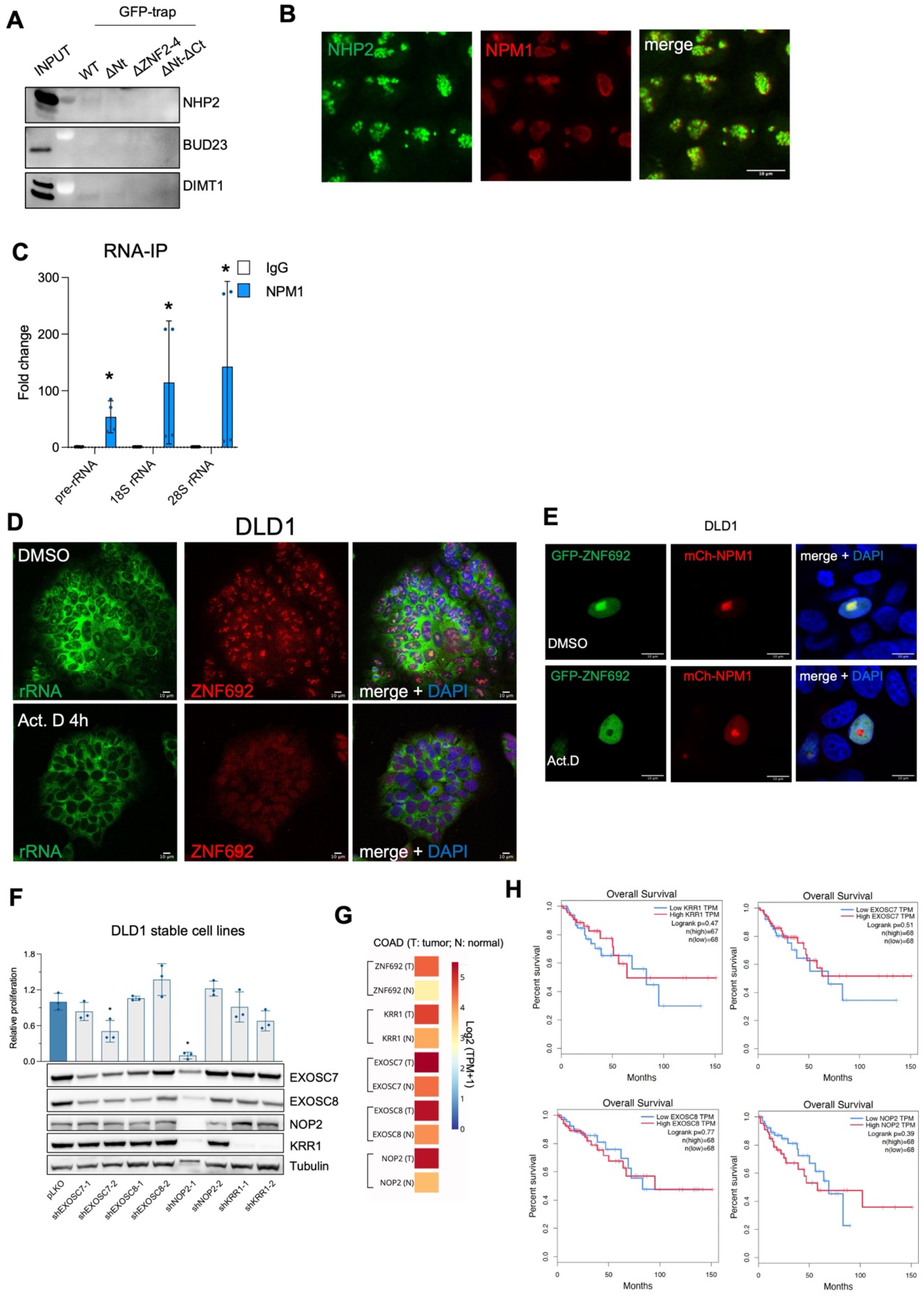
ZNF692 interacts with the components of the exosome complex. (A) IP of the recombinant ZNF692 using GFP nanotrap beads in the presence of nuclear extracts of DLD1 cells followed by immunoblot for NHP2, BUD23 and DIMT1 showing that these proteins did not co-IP with ZNF692 according to WB. (B) IF of NHP2 together with NPM1 showing that NHP2 and NPM1 do not colocalized. NPM1 is present in the granular component while NHP2 localizes in the DFC of the nucleolus. (C) RNA**-**IP of NPM1 on rRNA in DLD1 cells. NPM1 binds rRNA in DLD1 cells. Two independent IPs with two technical replicates each are represented. (D) IF of rRNA and ZNF692 in DLD1 cells treated with 0.05 μg/ml Actinomycin D for 4h. (E) IF of DLD1 cells transfected with GFP-ZNF692 and mCherry-NPM1 constructs and treated 4 h with DMSO or 0.05 μg/ml of Actinomycin D (Act. D). (F) Proliferation of DLD1 stably expressing shRNA for either control or NOP2, KRR1, EXOSC7, or EXOSC8. (G) Heatmap showing the expression of ZNF692, KRR1, EXOSC7, EXOSC8, and NOP2 in tumor vs normal tissues of COAD patients deposited in TCGA. (H) Patient survival correlation in relation with KRR1, EXOSC7, EXOSC8, and NOP2 mRNA levels in COAD patients from TCGA.

**Supplementary Figure 6:**
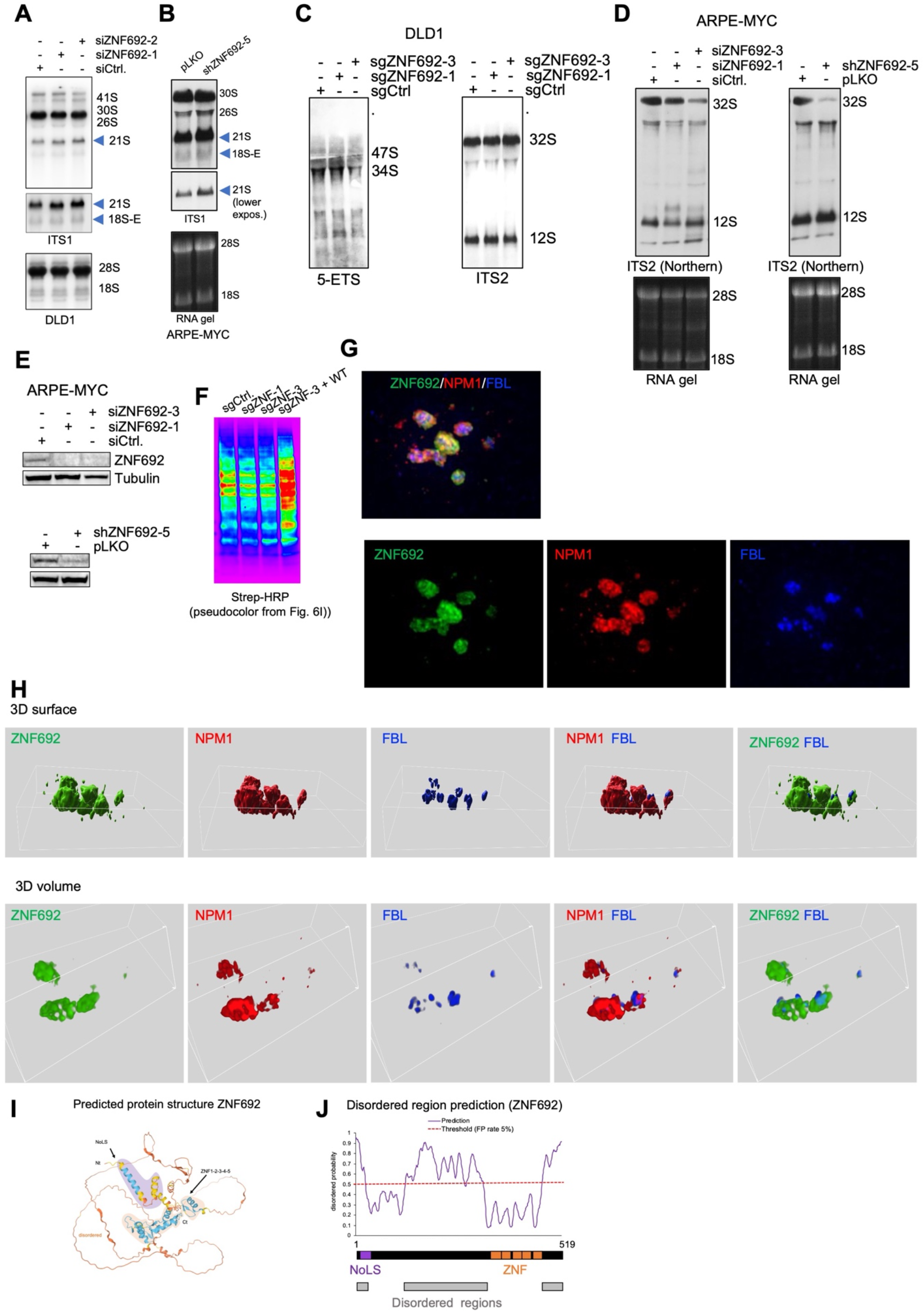
ZNF692 enhances 18S rRNA processing and small ribosomal subunit assembly. (A) Northern blot of DLD1 cells transiently transfected with siRNA for ZNF692 or control. Blue arrows indicate the 21S and 18S-E fragments. (B) Northern blot of ARPE cells stably overexpressing MYC and transiently infected with lentiviral particles containing shRNA for ZNF692 or control. Blue arrows indicate the 21S and 18S-E fragments. (C) Northern blot of control or ZNF692 CRIPSR KO DLD1 cells using 5’ETS (D) and ITS2 (E) probes. Mapping of the probes are shown in Fig. 6A. (D) Northern blot for ITS2 in ARPE cells stably overexpressing MYC and transiently transfected with siRNA for ZNF692 or control or infected with shRNA for ZNF692 or control (lower panel). Mapping of the probes are shown in Fig. 6A. (E) WB for ZNF692 in ARPE cells stably overexpressing MYC and transiently transfected with siRNA for ZNF692 or control (upper panel) or infected with shRNA for ZNF692 or control (lower panel). (F) Click-IT^TM^ AHA experiment from Fig. 6I showing pseudo colors, red indicates more intense bands while blue less intense. (G) IF of ZNF692, NPM1 and FBL in DLD1 cells overexpressing ZNF692. Image was acquired in Z-stack and is shown as a projection of the planes. 3D reconstruction of the images in (M) showing the surface (left panel) and the volume (right panel) of ZNF692, NPM1, and FBL in the nucleolus. (H) 3D reconstruction of immunofluorescence images (Fig. S6F) for NPM1, ZNF692 and FBL in DLD1 KO cells stably overexpressing ZNF692 showing the surface (upper panels) and the volume (lower panels) of ZNF692, NPM1, and FBL in the nucleolus. (I) ZNF692 predicted protein structure using AlphaFold showing that ZNF692 contains a structured N and C terminus region linked by a disordered central domain. (J) Plot and schematic representation of ZNF692 protein sequence showing the NoLS, Zn fingers, and the predicted disordered regions on ZNF692 identified with Protein DisOrder prediction System (PrDOS). ZNF692 central domain contain a large, disordered region.

**Supplementary Table 1.**
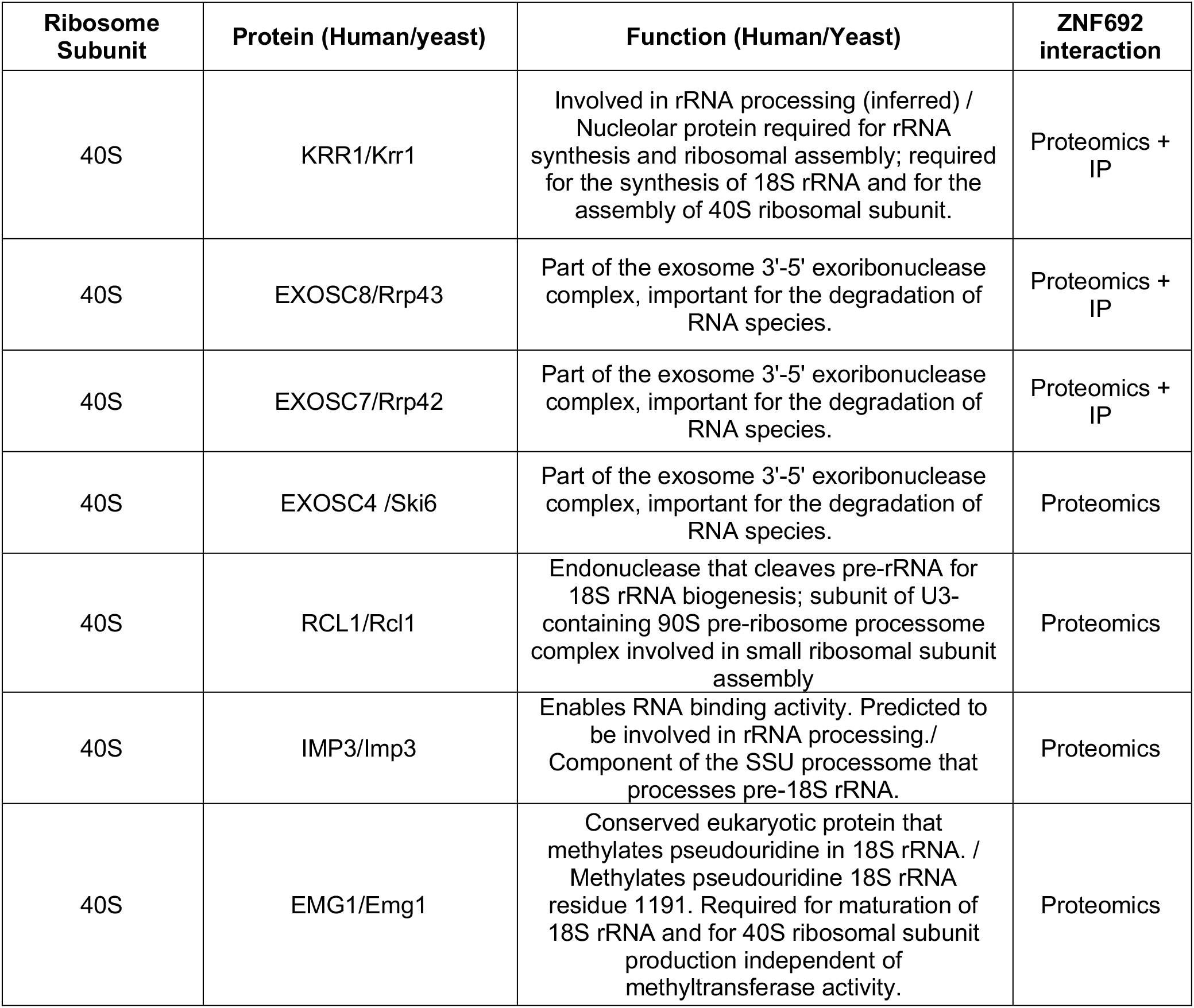
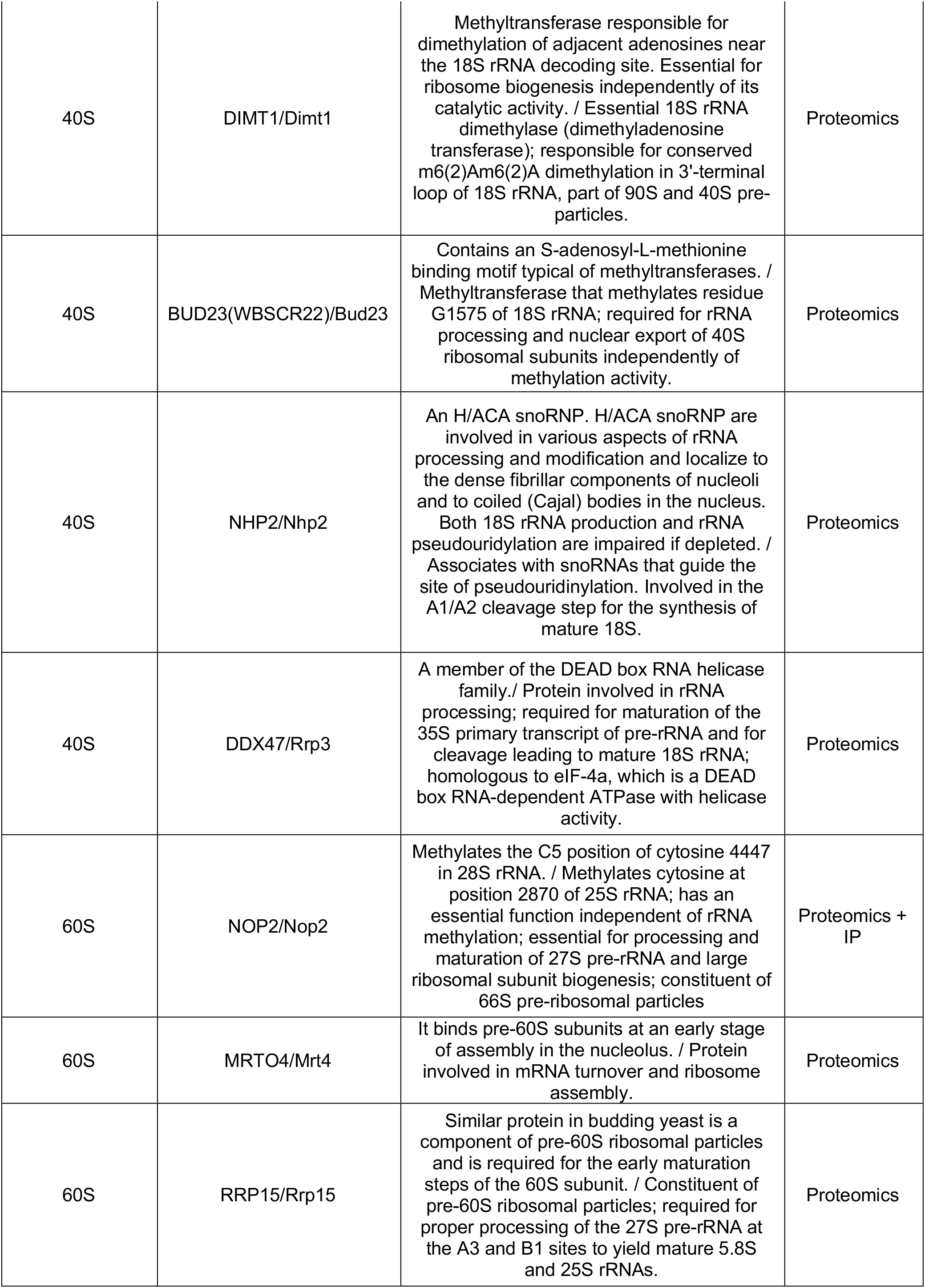
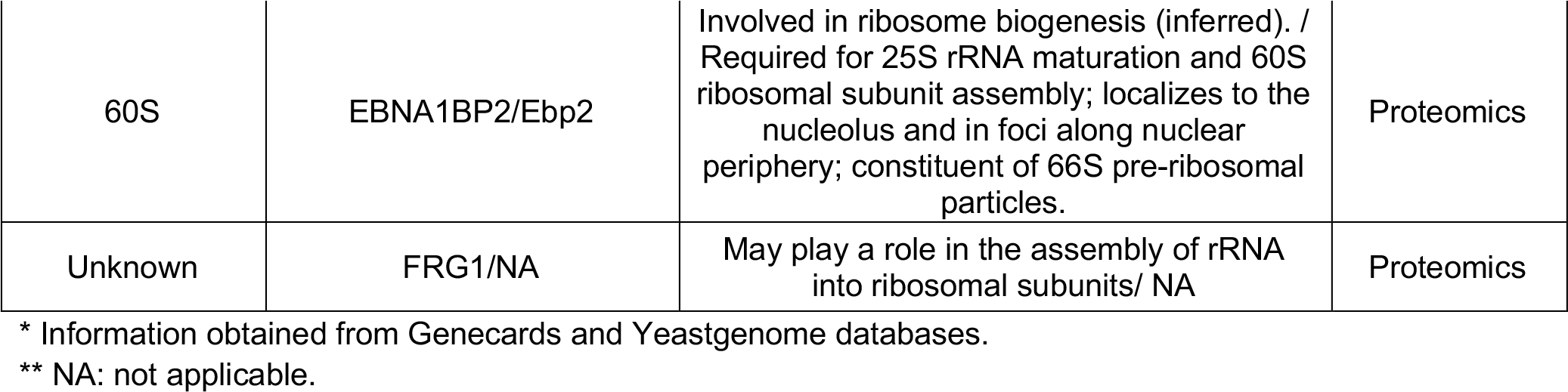
ZNF692 interactors found by IP-proteomics. List of ZNF692 interactors found in this work.

**Supplementary Table 2.**
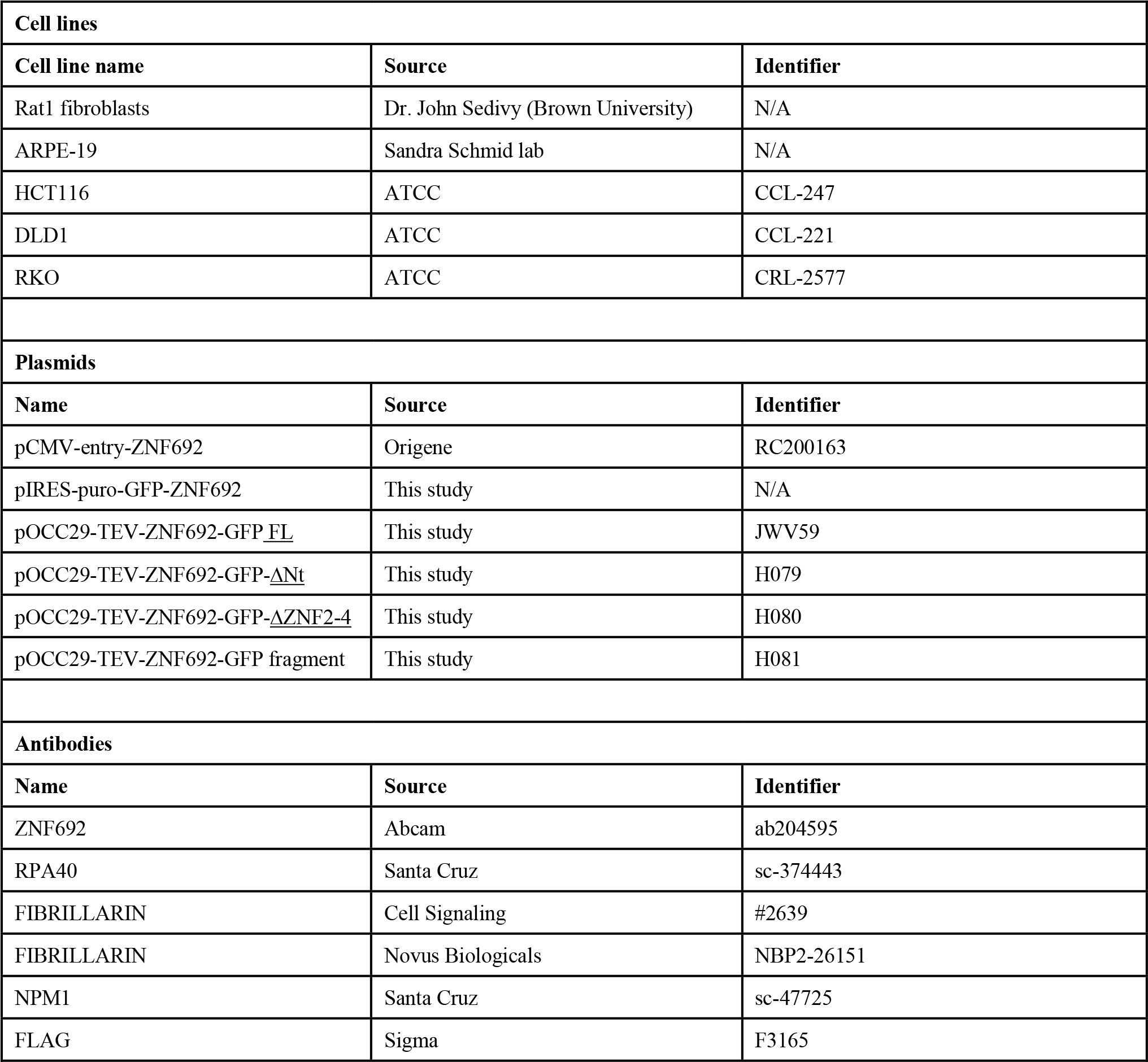
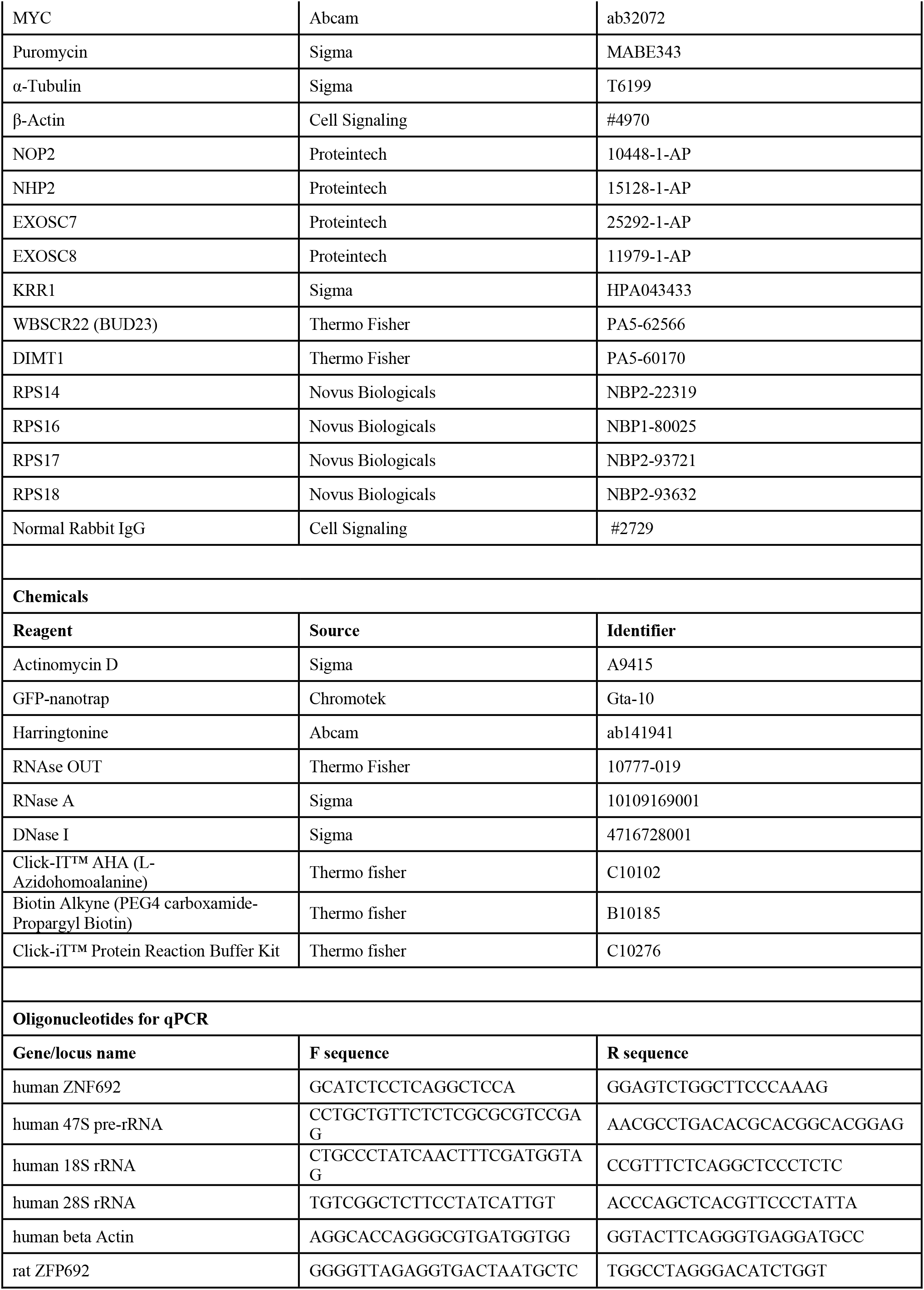
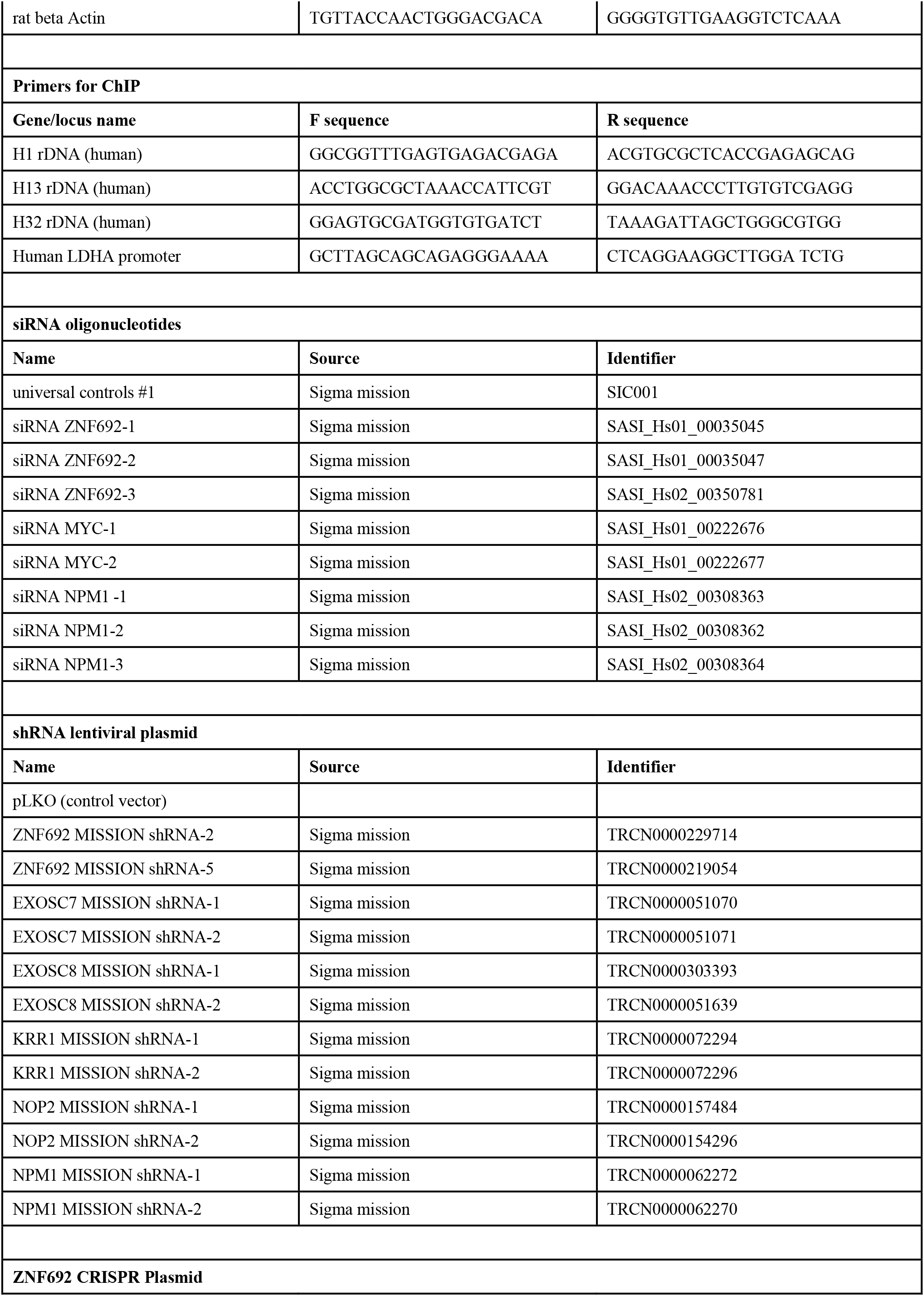
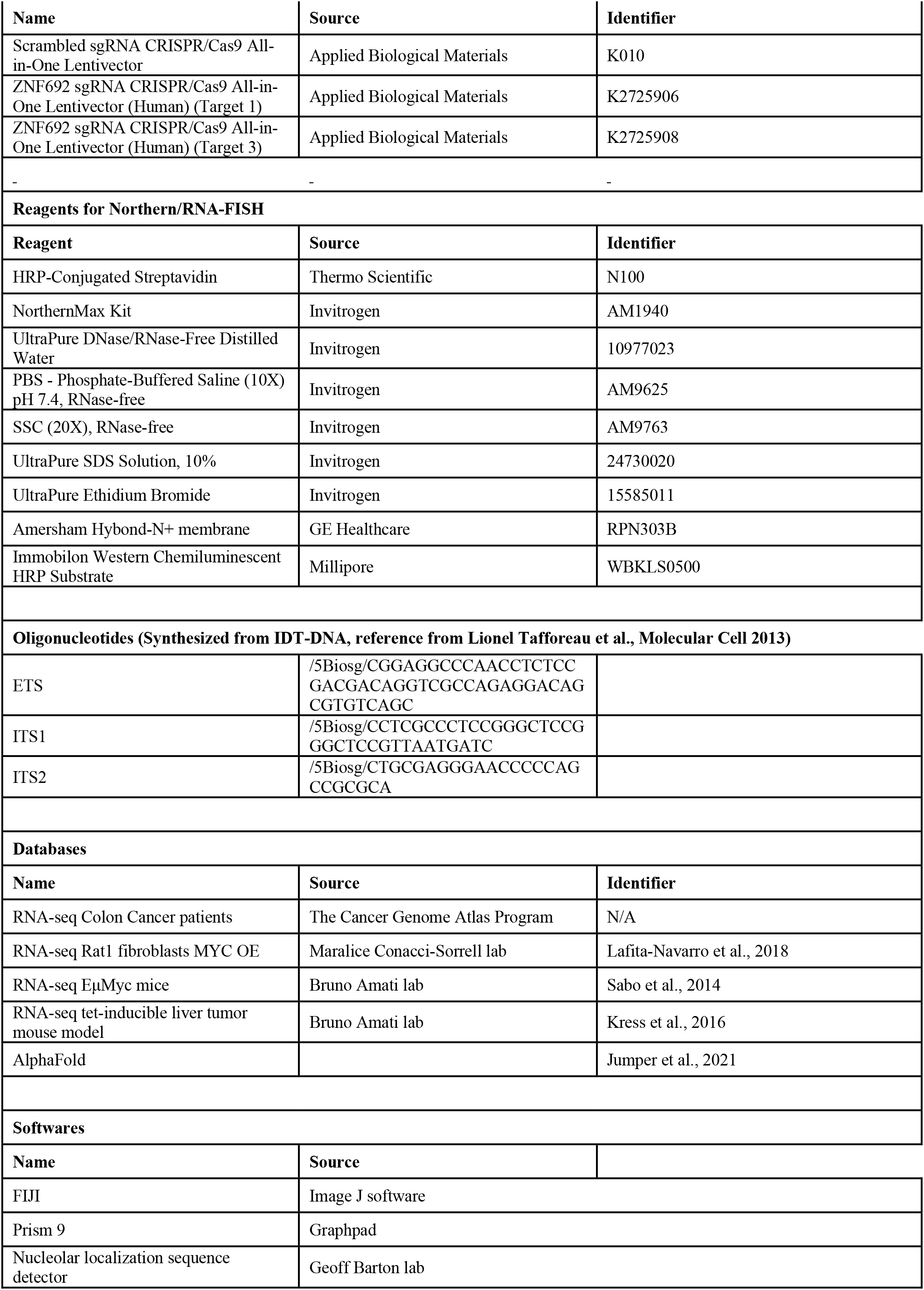
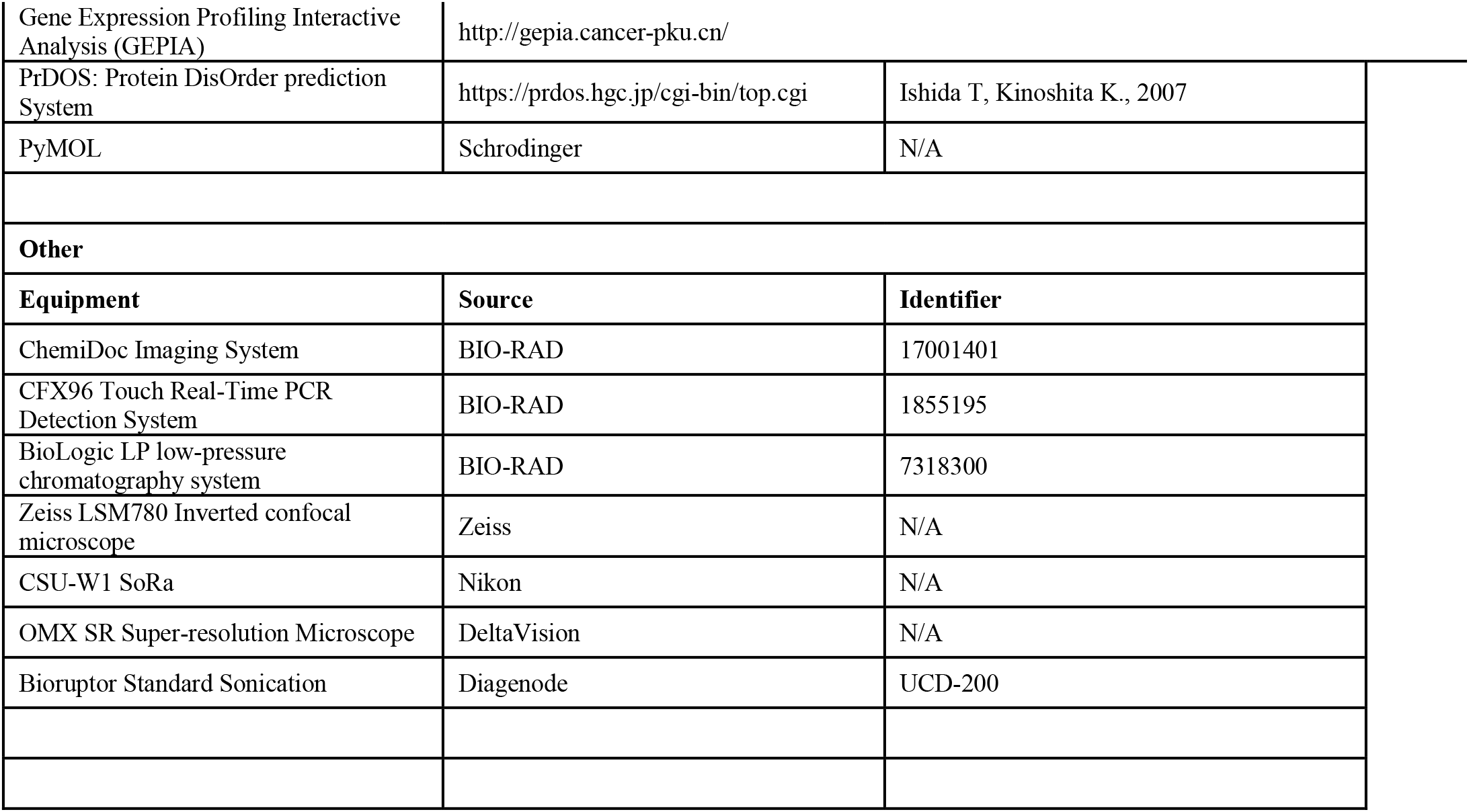
Reagents used in this work. Reagents, softwares and databases used in the present work are listed in this table.

## Methods

### Cell culture

Rat1 fibroblasts, human epithelial cell line ARPE-19, and human colon cancer cell lines (HCT116 and DLD1) were cultured in Dulbecco’s Modified Eagle Medium (DMEM) supplemented with 5% Fetal Bovine Serum (FBS), 100 U/mL Penicillin-Streptomycin.

### Plasmids

The pCMV6-Entry-ZNF692 plasmid was purchased from Origene. Human cDNAs of full length ZNF692 were amplified by PCR and cloned to pIRES-puro-GFP and pIRES-puro-FLAG vectors with restriction enzymes FseI and AscI. For *in vitro* purified protein, cDNAs of ZNF692 and NPM1 that were optimized for baculovirus expression were synthesized and cloned to pOCC29-TEV-GFP and pOCC206-mCherry vectors, respectively, with restriction enzymes NotI and AscI. To make the plasmids expressing fragment of ZNF692, corresponding cDNA sequence were amplified from full length plasmids and cloned using same pairs of restriction enzymes. Deletion mutations of ZNF692 plasmids were generated by PCR using Pfu polymerase.

### Transient Transfections

siRNA was reverse transfected to cells with Lipofectamine RNAiMAX reagent following the standard protocol (Invitrogen). Briefly, on the day of transfection, 1×10^5^ cells were seeded per well in 12-well plates and transfected with 2 µL siRNA (20 µM) and 2 µL Lipofectamine previously mixed in OPTI-MEM for 15 min. Cells were collected and analyzed 72hs after transfection.

ZNF692 overexpression was performed with Lipofectamine 3000 reagents following the standard protocol (Invitrogen). Briefly, one day before transfection, 1×10^6^ cells were seeded per well in 6-well plates. On the day of transfection, 2.5 µg DNA plasmid and 5ul Lipofectamine reagent were mixed and incubated for 15 min, then added to cells. Cells were collected for analysis after 72hs transfection. siRNAs used in this work are listed in Supplementary Table 2.

### Viral transduction

100,000-200,000 cells were seeded and mixed with 10 µg/ml polybrene and pLKO (lentiviral empty vector) or shRNA containing lentivirus made in 293T cells. For in transient experiments, after the indicated times, cells were harvested for western blot analysis or stained with crystal violet to assess proliferation.

To make stable cell lines, 48 h after adding the lentiviral particles, media was replaced and puromycin was added to select the cells containing the pLKO (lentiviral empty vector) or shRNA constructs. 10 µg/ml puromycin was used for DLD1 and 5 µg/ml puromycin was used for HCT116 cells. See Supplementary Table 2. Plasmids used in this work are listed in Supplementary Table 2.

### ZNF692 CRISPR knockout cell lines

ZNF692 sgRNA CRISPR/Cas9 All-in-One Lentivector or Scrambled sgRNA CRISPR/Cas9 All-in-One Lentivector (Applied Biological Materials) was transfected together with lentiviral packaging plasmids pSPAX2 and pMD2g to HEK293FT cells. The conditional media of transfected cells (viruses) were filtered with 0.45 µm filter and used to infect DLD1 cells. Knockout cells were selected with 10 µg/ml puromycin. Plasmids used in this work are listed in Supplementary Table 2.

### RT-qPCR

Total RNA was extracted with Tri-Reagent (sigma) following manufacturer’s instructions. RNA was reverse transcribed to cDNA with High-Capacity cDNA Reverse Transcription Kit (Life Technologies). RNA levels were measured by quantitative PCR with the iTaq™ Universal SYBR® Green Supermix (BIO-RAD) and the BioRad CFX96 device. For qPCR result analysis, 2^−ΔΔCt^ method was used. qPCR primers are listed in key resources table. All RT-qPCR reactions were normalized by Actin mRNA levels (used as housekeeping gene). Primers used in this work are listed in Supplementary Table 2.

### Western Blot

Total protein was extracted with lysis buffer containing 10 mM Tris-HCl (pH 8.0), 50 mM NaCl, 0.1% Nonidet P-40 or RIPA buffer (25 mM Tris-HCl pH 7.4, 150 mM NaCl, 1% NP-40, 0.5 % sodium deoxycholate, 0.1 % SDS) + protease and phosphatase inhibitors and MG132. Protein concentrations were measured by Pierce BCA protein assay. Proteins were separated by SDS–polyacrylamide gel electrophoresis, then transferred to nitrocellulose membranes (Thermo Fisher), and probed with specific antibodies (See Key Resource Table), then detected by chemiluminescence with BioRad ChemiDocImager system. Antibodies used in this work are listed in Supplementary Table 2.

### Immunofluorescence staining and microscopy imaging

Cells were grown on glass coverslips and fixed with 4% paraformaldehyde PBS for 15 min, permeabilized with 0.1 % Triton X-100 for 20 min and blocked with 5% BSA in PBS for 30 min-1 h. Primary antibodies were diluted in 5% BSA-PBS and incubated overnight at 4°C. The cells were washed 3 times for 10 min PBS and incubated with secondary antibodies. Secondary antibodies were diluted in 5% BSA-PBS, incubated for 1 h at room temperature, washed with PBS 3 times for 10 min and mounted with Mowiol mounting media. The second PBS wash contained DAPI at 5 µg/mL to stain the nuclei. Images were acquired with a Zeiss LSM780 or Nikon CSU-W1 SoRa fluorescent microscope. For SIM images (Fig.2J), DeltaVision OMX SR Super-resolution Microscope was used (GE Healthcare).

The antibodies used in this study are indicated in the Supplementary Table 2. Images were processed with FIJI.

### In silico analyses

The online software Nucleolar localization sequence Detector, Protein DisOrder prediction System and AlphaFold were used to predict the presence of nucleolar localization sequences, disordered regions, and the protein structures by providing human ZNF692 and NPM1 protein sequences.

For ribosome reconstructions, the structure of the human 80S ribosome (PDB: 4V6X) was obtained from Protein Data Bank (https://www.rcsb.org/). Specific ribosomal proteins found to interact with ZNF692 by IP-proteomics analysis were highlighted with PyMOL software.

### TCGA Data Analysis

RNA expression and patient survival analysis from the RNA-seq experiments deposited in The Cancer Genome Atlas (TCGA) was performed with the Gene Expression Profiling Interactive Analysis (GEPIA) web server http://gepia.cancer-pku.cn/.

### Electron microscopy

HCT116 cells were transiently infected with lentiviral particles containing control (pLKO) or shRNA for ZNF692. After 3 days of infection cells were fixed on MatTek dishes with 2.5% (v/v) glutaraldehyde in 0.1M sodium cacodylate buffer. After three rinses in 0.1 M sodium cacodylate buffer, they were post-fixed in 1% osmium tetroxide and 0.8 % K3[Fe(CN6)] in 0.1 M sodium cacodylate buffer for 1 h at room temperature. Samples were rinsed with water and en bloc stained with 2% aqueous uranyl acetate overnight. After three rinses with water, they were dehydrated with increasing concentration of ethanol, infiltrated with Embed-812 resin and polymerized in a 60oC oven overnight. Blocks were sectioned with a diamond knife (Diatome) on a Leica Ultracut UCT (7) ultramicrotome (Leica Microsystems) and collected onto copper grids, post stained with 2% Uranyl acetate in water and lead citrate. Images were acquired on a JEOL 1400 Plus (JEOL) equipped with a LaB6 source using a voltage of 120 kV.

To perform the nucleolar perimeter, and circularity measurements, FIJI was used.

### Puromycilation

Puromycin was added at 20 µg/ml for DLD1 and 10 µg/ml for HCT116. For the majority of the experiments, cells were serum starved overnight and the next day complete media (containing FBS) was added for 6 hours, then puromycin was added for 2 additional hours. For Fig. 5G, cells were seeded and the next day fresh media with puromycin was added for 20 min. After puromycin incubation, cells were harvested to lyse with RIPA buffer and perform Western blotting.

For harrigtonine experiments, cells were seeded in 6 well plates. The next day, harringtonine was added for 1, 2, and 8 min. After incubation with harringtonine, puromycin was added for additional 20 min, then cells were harvested to lyse with RIPA buffer and perform Western blotting.

The puromycin antibody used for this assay is listed in supplementary Table 2.

### Cytoplasmic isolation of single 40S, 60S and 80S ribosomes

Isolation of 40S, 60S and 80S ribosomes was performed by centrifuging cytoplasmic fraction over a sucrose gradient column. Details of experiment are indicated below.

#### Samples preparation

DLD1 control or CRISPR KO cells were used for these experiments. 4-5 million cells at 60%-80% confluency were harvested and resuspended in 500 µL lysis buffer (20 mM Tris pH 7.4, 5 mM MgCl_2_, 100 mM NaCl in DPEC-treated dH_2_O + 100 µg/mL CHX+ RNAse inhibitor RNAse OUT + Protease inhibitor cocktail (1/100 from stock) + 0.1% NP-40). The lysates were incubated on ice for 15 min resuspending every 5 min. Samples were centrifuge 12,000g 4℃ 10 min to pellet nuclei and mitochondria. Supernatants were collected. RNA amounts were quantified by nanodrop, and all samples were set at the same RNA concentration to load same amount of RNA in sucrose gradient columns.

#### Sucrose gradient preparation

Gradients were made by solubilizing different sucrose amounts in buffer 0 % (20 mM Tris pH 7.4, 5 mM MgCl_2_, 100 mM NaCl in DPEC-treated dH_2_O + 100 µg/mL CHX + RNAse inhibitor RNAse OUT). The sucrose gradient column consists of 2 mL of 10 %, 20 %, 30%, 40 %, and 50 % sucrose solutions. Starting the gradient column by the most concentrated sucrose solution. Each time a different gradient solution was added, the solution was frozen at −80 °C for at least 20 min before applying the next sucrose % solution. The columns were kept at −80 °C until use. Before use, the gradient column tubes were thawed at 4°C overnight to form continuous sucrose gradient.

#### Polysome fractionation to isolate single 40S, 60S and 80S ribosomes

Lysates were loaded on gradient columns on ice and vertical. The column walls were cleaned to remove any liquid before balancing, weighed, and balanced with <0.005 g difference of weight by adding buffer 0%. Balanced columns were centrifuged in a swinging bucket rotor at 34,000 rpm for 2 h at 4℃, acc=8, dec=0.

#### Polysome analysis to isolate single 40S, 60S and 80S ribosomes

The BioLogic LP low-pressure chromatography system (BIO-RAD) was used to analyze the centrifuged fractions and collect the fractions. The samples were run at 1 mL/min, and the UV recorded. Fractions were collected every 30 sec (0.5 ml fractions). Fractions corresponding to single 40S, 60S and 80S ribosomes according to the polysome profile were used for proteomics, WB, as well as RNA tapestation analysis to confirm the purity of the fractions.

This experiment was performed twice with similar results.

### Cell proliferation

Six-well plates were seeded with 1.5 x 10^5^ cells, while twelve-well plates were seeded with 50,000 cells (for 3-day experiments) or 20,000 cells (for 7-day experiments). Cell proliferation assays were performed with crystal violet staining. Briefly, cells were washed with PBS on plates, and fixed with methanol at room temperature for 10 min. Then cells were stained with crystal violet solution containing 1% acetic acid, 1% methanol, 1% (w:v) crystal violet dye for 10 min with agitation. After washing extensively, the crystal violet was extracted with 10 % glacial acetic acid and the absorbance was read at 595 nm. Results are presented to reflect the relative growth after normalization by the control condition.

### Xenografts experiments

For the xenograft experiment 1 x 10^6^ DLD1 control or ZNF692 CRISPR KO cells (sg1 and sg3) were injected into the flank of NOD/SCID mice (from Jackson lab). Tumors were measured once every week and animals were sacrificed when tumors reached 2 cm in volume. At the end of the experiment the tumors were harvested, weighed and snap frozen in liquid nitrogen and processed for protein and RNA extraction. All procedures are approved by IACUC, UT southwestern medical center.

### Nuclear extracts for *in vitro* protein immunoprecipitation (IP)

DLD1 nuclear extracts were obtained by lysing the cells with lysis buffer A (10 mM Tris-HCl (pH 8.0), 50 mM NaCl, 0.1% Nonidet P-40) + proteinase inhibitors for 20 min on ice and pipetting up and down every 5 min. After incubation, cells were centrifuge 1200 rpm for 5 min. Supernatants (cytoplasmic fractions) were discarded and nuclear pellets were lysed with buffer A + proteinase inhibitors 20 min and sonicated at max intensity (30 sec ON, 30 sec OFF) for 5 min. Then lysates were centrifuge at 15,000 g at 4 °C. Supernatant were collected as nuclear extracts. Antibodies used in this work are listed in Supplementary Table 2.

### Protein Immunoprecipitation (IP)

Cells were collected and lysed with lysis buffer A containing 10 mM Tris-HCl (pH 8.0), 50 mM NaCl, 0.1% Nonidet P-40, and proteinase inhibitor for 20 mins on ice. Then, the samples were sonicated and centrifuged at 15,000g at 4 °C for 20 min to remove the debris.

Same amount of protein lysate was use for IP and 10% of each sample was saved for input. Lysates were incubated with the primary antibodies or GFP nanotrap magnetic beads rotating overnight at 4°C.

For IP with primary antibodies, magnetic beads were added to the lysate-antibody mixture for 3 additional h rotating at 4°C.

For the IP with the GFP-ZNF692 purified proteins, 5 µM of each construct was incubated with GFP nanotrap magnetic beads and equal amounts of DLD1 nuclear extracts overnight at 4°C.

After incubation, beads containing the immunocomplexes were washed with lysis buffer A at least 3 times for 10 min. Immunoprecipitates were eluted by incubating the antibody-beads complexes with 2 × SDS Laemli sample buffer and boiling them for at least 15 min (vortexing every 5 min). Samples were spined down and beads removed by using a magnet. Supernatants were collected and subjected to western blot analysis.

Antibodies and reagents used in this work are listed in Supplementary Table 2.

### Proteomics

HCT116 were transfected with GFP, GFP-ZNF692 or GFP-ZNF692 ΔNoLS plasmids. Cells were collected and lysed with lysis buffer A containing 10 mM Tris-HCl (pH 8.0), 50 mM NaCl, 0.1% Nonidet P-40, and proteinase inhibitor for 20 mins on ice. Then, the samples were sonicated and centrifuged at 15,000g at 4 °C for 20 min to remove the debris. Supernatants were used for IP with GFP nanotrap magnetic beads. Eluted lysates were ran on gel and proteins were extracted for proteomics analysis at UTSW proteomics core.

To identify ZNF692’s interactome first a stringent fold change relative to GFP alone was applied (Fold change WT vs GFP >10, and fold change ΔNt vs GFP >10). Then, to identify nucleolar ZNF692 binding partners, we only considered the interactors that lost their binding to ZNF692 when ZNF692 nucleolar localization signal was deleted. The levels of immunoprecipitated ZNF692 with GFP-trap beads was about 4 times higher in the WT than the ΔNt-ZNF692 (FigS4B). Thus, we a cutoff relative to the amounts of ZNF692 found in the proteomics for WT or ΔNt was applied to find the partner specifically dependent on the NoLS region of ZNF692 (fold change WT_vs_GFP vs ΔNt_vs_GFP > or = 4). To identify ZNF692 interactors that were not dependent on the NoLS a fold change WT_vs_GFP vs ΔNt_vs_GFP < 4 was applied.

### ChIP-qPCR

Cells at 70% confluency were fixed with 1 % formaldehyde. Cells were lysed with lysis buffer A (10 mM Tris-HCl (pH 8.0), 50 mM NaCl, 0.1% Nonidet P-40) + proteinase inhibitors for 20 min on ice and pipetting up and down every 5 min. After incubation, cells were centrifuge 1200 rpm for 5 min. Supernatants (cytoplasmic fractions) were discarded and nuclear pellets were lysed with RIPA buffer for 10 min on ice and pipetting up and down every 5 min. DNA was sonicated to obtain ∼500 bp fragments using a the diagenode sonicator at high intensity (30 sec ON, 30 sec OFF). After sonication, cells were centrifuged 15,000 rpm for 15 min at 4 °C. Supernatants were collected as nuclear lysates. Nuclei lysates were diluted 1:10 for immunoprecipitation with ZNF692 or POLR1 antibody. Normal rabbit IgG was used a negative control. Next, beads were washed with lysis buffer A 3 times. To reverse DNA crosslinking, beads were resuspended with 200 ul of elution buffer (1 % SDS; 50 mM Tris-HCl pH 8) and incubated with 12 µl of 10 mg/ml RNAse A and 24 µl 5M NaCl overnight at 65 °C in agitation making sure the magnetic beads do not precipitate. The next day add 4 µl of 10 mg/ml proteinase K, 4 μl 0.5 M EDTA and 8 μl 1M Tris-HCl pH 6.8 was added to the samples and they were incubated for at least 3 h at 45 °C in agitation to degrade proteins were added to degrade the protein in the mixture. After incubation, beads were removed with the magnet and DNA purified from the supernatants for DNA purification and analysis by qPCR. Fold change enrichment was normalized by comparing the amount of DNA immunoprecipitated with the specific antibodies in comparison with normal IgG. The primers used in this study are indicated in supplementary Table 2.

### RNA-IP

Cells at 70% confluency were crosslinked with UV crosslink on ice in a Spectrolinker XL-1500 at 254nm at 400mJ/cm^2^. Cells were lysed with lysis buffer A (10 mM Tris-HCl (pH 8.0), 50 mM NaCl, 0.1% Nonidet P-40) + proteinase and RNAse inhibitors for 20 min on ice and pipetting up and down every 5 min. After incubation, cells were centrifuge 1200 rpm for 5 min. Supernatants (cytoplasmic fractions) were discarded and nuclear pellets were lysed with RNA-IP lysis buffer (50mM Tris-HCl pH 7.4, 100mM NaCl, 1% Nonidet P-40, 0.1% SDS, 0.5% sodium deoxycholate (protect from light)) + proteinase and RNAse inhibitors for 20 min on ice and pipetting up and down every 5 min. Then, cells were centrifuged 15,000 rpm for 15 min at 4 °C. Supernatants were collected as nuclear lysates. Same amount of nuclei lysates were used for immunoprecipitation. First, magnetic beads washed with lysis buffer were incubated with no antibody (as negative control) or antibodies for ZNF692 and NPM1 for 45 min at room temperature. Next, nuclear lysates were added to the antibody-bead complexes and incubated over night at 4 °C. Then, beads were washed 3 times with High Salt Wash Buffer (50mM Tris-HCl pH 7.4, 1 M NaCl, 1mM EDTA, 1% NP-40, 0.1% SDS, 0.5% sodium deoxycholate (protect from light)) and 3 times with Wash Buffer #2 (20mM Tris-HCl pH 7.4, 10mM MgCl2, 0.2% Tween-20). After washes, beads were resuspended in 100 µl of wash buffer #2. DNAse I was added to the mixture to degrade DNA and incubated for 1-2 h at 37 °C in agitation. Then, 500 µl of TRI-Reagent was added to the mixture followed by 100 µl chloroform. The mixed was incubated at room temperature for 5 min and then centrifuge for 10 min at 12,000 rpm. The upper transparent phase was then collected and mixed with 1x volume of 70% ethanol. To continue with RNA purification RNeasy Kit from Qiagen was used. To quantify the amount of RNA immunoprecipitated, half of the RNA eluted from the column was reverse transcribed to cDNA with High-Capacity cDNA Reverse Transcription Kit (Life Technologies) while the other half was incubated with the buffer containing no retro-transcriptase as a negative control. After cDNA synthesis, quantitative PCR was performed with the iTaq™ Universal SYBR® Green Supermix (BIO-RAD) and the BioRad CFX96 device. For qPCR result analysis, 2^−ΔΔCt^ method was used. After that, the amount of RNA in each condition was first normalized by the amount of that condition without retro-transcriptase to account for unspecific quantification. Then each condition compared with the no antibody conditions to determine the fold change enrichment. The primers used in this study are indicated in supplementary Table 2.

### Northern

DLD1 sgCtrl., sgZNF691-1, and sgZNF692-3 cells were seeded in 10 cm dishes. Then cells were starved ON and fresh media containing FBS was added the following morning for 24 h. Then, cells were harvest for RNA extraction with TriZol.

ARPE-MYC were transiently infected with lentiviral particles containing control (pLKO) or shRNA for ZNF692. After 3 days of infection cells were starved ON and fresh media containing FBS was added the following morning for 24 h. Then, cells were harvest for RNA extraction with TriZol.

ARPE-MYC cells were seeded in 6 cm dishes and transfected with 10 µL control siRNA or siRNAs against ZNF692 (20 µM) and 10 µL Lipofectamine previously mixed in OPTI-MEM for 15 min. 2 days after infection cells were starved ON and fresh media containing FBS was added the following morning for 24 h. Then, cells were harvest for RNA extraction with TriZol. Purified RNA was mixed with 3 volumes of Formaldehyde Load Dye (Northern Max Kit, Invitrogen) containing 0.2 mg/mL Ethidium Bromide (Invitrogen) and incubated at 55℃ for 15 min and then placed to ice for 2 min. 7.5-10 μg of total RNA were loaded onto a 1% agarose denaturing gel (Northern Max Kit, Invitrogen) and electrophoresed at 90 V for 3 hr. RNAs in the gel were then visualized using a UV gel imager. Next, RNAs were transferred to a positively charged nylon membrane (GE Healthcare) for 3 hr. RNAs on the membrane were crosslinked using UV light (120000 μJoules x three pulses) and then visualized using a UV gel imager. Membranes were prehybridized with ULTRAhyb buffer (Northern Max Kit, Invitrogen) for 30 min at 37℃ and hybridized overnight with each probe (10 μM, 1:1000) in ULTRAhyb buffer at 37℃. After washing with Stringency wash buffer (2X SSC and 0.5% SDS) twice and Wash buffer (0.5% SDS and 1X PBS) twice, membranes were blocked with Blocking buffer (0.5% SDS, 1% BSA and 1X PBS) for 30 min, followed by incubation with HRP-Conjugated streptavidin (Thermo Scientific, 1:4000) for 1 hr at 37℃. The membranes were then washed with Wash buffer three times and 1X PBS three times. The RNA bands were detected with Immobilon Western Chemiluminescent HRP Substrate (Millipore).

For experiment in Fig. S6A, HCT116 cells were transiently transfected with siCtrl. or siZNF692, after 3 days of transfection, cells were harvest for RNA extraction.

For this experiment, northern was performed as follows: 10 µg RNA was mixed with Sample Loading Buffer (Sigma, cat. #R4268), heated at 65°C for 10 minutes and then placed to ice. Samples were loaded onto an agarose gel and electrophoresed at 85 V for 360 Vh. Next, RNAs were transferred to GeneScreen Plus Membrane (Perkin Elmer, cat. #NEF976) overnight, crosslinked in a UV Crosslinker for 50mJ, and baked 1 and a half hours at 80°C in vacuum oven. Membranes were then prehybridized with ULTRAhyb oligo buffer (Ambion Cat #:8663) for 30 min at 37℃ and hybridized overnight at 42℃ with probe that was labeled with γ-^32^P-ATP (Perkin Elmer Cat # NEG035C; 6000 Ci/mmol). Expose as needed after washing with 2X SSC and 0.5% SDS for 3 times.

### RNA-FISH

Cells were grown on glass coverslips and fixed with 4% paraformaldehyde PBS for 15 min, permeabilized with 0.1 % Triton X-100 for 20 min. Coverslips were rinsed once with room temperature PBS and then hybridized with NorthernMax™ Prehybridization/Hybridization Buffer (Thermo Fisher) in a humid chamber for 1 h at 65°C. Then prehybridization buffer was removed. 20 µL of Hybridization buffer containing 55 nM biotin-labeled probes 5’ETS (indicated in supplementary Table 2) were added overnight at 65°C in a humid chamber. Then coverslips were washed three times with 2× SSC at 37°C and twice t with 1× SSC at room temperature. Slides were then refixed in 4% formaldehyde PBS for 15 min at c, washed twice with cold 1× PBS and blocked with PBS-BSA for 30 min at room temperature. Streptavidin-Alexa 568 was used at 1:200 dilution for 1 h at room temperature in the dark. Coverslips were then washing twice with PBS, and one final washes with PBS+DAPI before mounting with mowiol. Coverslips were left ON at room temperature before imaging. Images were acquired with a Zeiss LSM780. Images were process using FiJi software.

### Quantification and statistical analysis

All statistical analyses were performed using two-tailed unpaired T-student statistical analysis, p ≦ 0.05 was considered statistically significant. All values are reported as mean ± SD in each figure.

